# A PTER-dependent pathway of taurine metabolism linked to energy balance

**DOI:** 10.1101/2024.03.21.586194

**Authors:** Wei Wei, Xuchao Lyu, Andrew L. Markhard, Sipei Fu, Rachel E. Mardjuki, Peter E. Cavanagh, Xianfeng Zeng, Jakub Rajniak, Nannan Lu, Shuke Xiao, Meng Zhao, Maria Dolores Moya-Garzon, Steven D. Truong, Jonathan Chiu-Chun Chou, Lianna W. Wat, Saranya Chidambaranathan-Reghupaty, Laetitia Coassolo, Duo Xu, Fangfang Shen, Wentao Huang, Cuauhtemoc B. Ramirez, Cholsoon Jang, Katrin J. Svensson, Michael A Fischbach, Jonathan Z. Long

## Abstract

Taurine is a conditionally essential micronutrient and one of the most abundant amino acids in humans^1–3^. In endogenous taurine metabolism, dedicated enzymes are involved in biosynthesis of taurine from cysteine as well as the downstream derivatization of taurine into secondary taurine metabolites^4,5^. One such taurine metabolite is N-acetyltaurine^6^. Levels of N-acetyltaurine are dynamically regulated by diverse physiologic perturbations that alter taurine and/or acetate flux, including endurance exercise^7^, nutritional taurine supplementation^8^, and alcohol consumption^6,9^. While taurine N-acetyltransferase activity has been previously detected in mammalian cells^6,7^, the molecular identity of this enzyme, and the physiologic relevance of N-acetyltaurine, have remained unknown. Here we show that the orphan body mass index-associated enzyme PTER (phosphotriesterase-related)^10^ is the principal mammalian taurine N-acetyltransferase/hydrolase. In vitro, recombinant PTER catalyzes bidirectional taurine N-acetylation with free acetate as well as the reverse N-acetyltaurine hydrolysis reaction. Genetic ablation of PTER in mice results in complete loss of tissue taurine N-acetyltransferase/hydrolysis activities and systemic elevation of N-acetyltaurine levels. Upon stimuli that increase taurine levels, PTER-KO mice exhibit lower body weight, reduced adiposity, and improved glucose homeostasis. These phenotypes are recapitulated by administration of N-acetyltaurine to wild-type mice. Lastly, the anorexigenic and anti-obesity effects of N-acetyltaurine require functional GFRAL receptors. Together, these data uncover enzymatic control of a previously enigmatic pathway of secondary taurine metabolism linked to energy balance.

## Introduction

Taurine is a conditionally essential micronutrient and very abundant amino sulfonic acid that is found in mammalian tissues and many foods^2,11^. Levels of taurine are especially high in excitable tissues such as the heart, eyes, brain, and muscles^5^. Taurine has been described to have pleiotropic cellular and physiologic functions, particularly in the context of metabolic homeostasis^11–13^. Genetic reduction of tissue taurine levels leads to muscle atrophy^14,15^, decreased exercise capacity^16^, and mitochondrial dysfunction in multiple tissues^14,17^. Conversely, taurine supplementation has been reported to reduce mitochondrial redox stress^11^, enhance exercise performance^18^ and suppress body weight^19^.

The biochemistry and enzymology of taurine metabolism has attracted considerable research interest. In the endogenous taurine biosynthesis pathway, cysteine is metabolized via CDO and CSAD to generate hypotaurine^20,21^, which is subsequently oxidized by FMO1 to produce taurine^22^. In addition, cysteine can undergo an alternative pathway via cysteamine and ADO^23^. Downstream of taurine itself are several secondary taurine metabolites that include taurocholate, taurocyamine, and N-acetyltaurine^4^. The only enzyme known to catalyze one of these downstream pathways is BAAT, which conjugates taurine to bile acyl-CoAs to produce taurocholate and other bile salts^24^. Beyond BAAT as the sole example, the molecular identities of the additional enzymes that mediate secondary taurine metabolism have not yet been established.

The biochemical interconversion of taurine and N-acetyltaurine is of particular interest for several reasons. First, N-acetyltaurine is an abundant endogenous metabolite whose levels are dynamically regulated by diverse physiologic perturbations that increase taurine and/or acetate flux, including endurance exercise^7,19^, alcohol consumption^6,9^, and nutritional taurine supplementation^19^. Second, N-acetyltaurine exhibits chemical structural similarities with signaling molecules including the neurotransmitter acetylcholine^25^ and the glucoregulatory long-chain N-fatty acyl taurines^26^. Third, a taurine N-acetyltransferase biochemical activity has been detected in cells^6,7^, demonstrating that the production of N-acetyltaurine is not simply a byproduct of taurine metabolism, but rather an enzymatically regulated biochemical transformation.

Using an activity-guided fractionation approach, here we identify PTER (phosphotriesterase-related), an orphan enzyme of previously unknown function, as the principal mammalian taurine N-acetyltransferase/hydrolase. In vitro, PTER catalyzes bidirectional taurine N-acetyltransferase using free acetate as an acyl donor as well as N-acetyltaurine hydrolysis. PTER exhibits a narrow substrate scope that is largely restricted to taurine, acetate, and N-acetyltaurine as substrates. Genetic ablation of PTER in mice abolishes tissue taurine N-acetyltransferase/hydrolase activities and results in concomitant elevation of N-acetyltaurine across tissues. Lastly, using genetic clues linking the human PTER locus to body mass index, we provide functional evidence that genetic ablation of PTER in mice, or pharmacological administration of N-acetyltaurine, suppresses body weight and adiposity. The full anorexigenic and anti-obesity effects of N-acetyltaurine require functional GFRAL receptors. These data define a PTER-dependent pathway of taurine metabolism linked to energy balance.

## Results

### Biochemical purification of a taurine N-acetyltransferase/hydrolase activity from kidney

To identify the enzyme(s) that mediates interconversion of taurine and N-acetyltaurine, we used an in vitro enzyme activity-guided approach to detect and purify a taurine N-acetyltransferase/hydrolase activity from mouse tissues (**Fig. 1a**). Total tissue homogenates were incubated with taurine (10 mM) and acetate (10 mM) and the formation of N-acetyltaurine was monitored by liquid chromatography-mass spectrometry (LC-MS). Most tissues had minimal or undetectable enzyme activity (**Fig. 1b**). By contrast, robust taurine N-acetyltransferase activity was observed in the kidney, and, to a lower extent, the liver and the quadriceps muscle (**Fig. 1b**). The observed activity in the kidney was higher using acetate as a substrate, rather than acetyl-CoA (**Fig. 1c**). Lastly, because the formation of the amide bond of N-acetyltaurine is reversible, we also tested tissue homogenates for the reverse N-acetyltaurine hydrolysis activity. The relative pattern of N-acetyltaurine hydrolysis activity across tissues was similar to that of N-acetyltaurine synthesis (**Fig. 1d**).

**Fig. 1.**
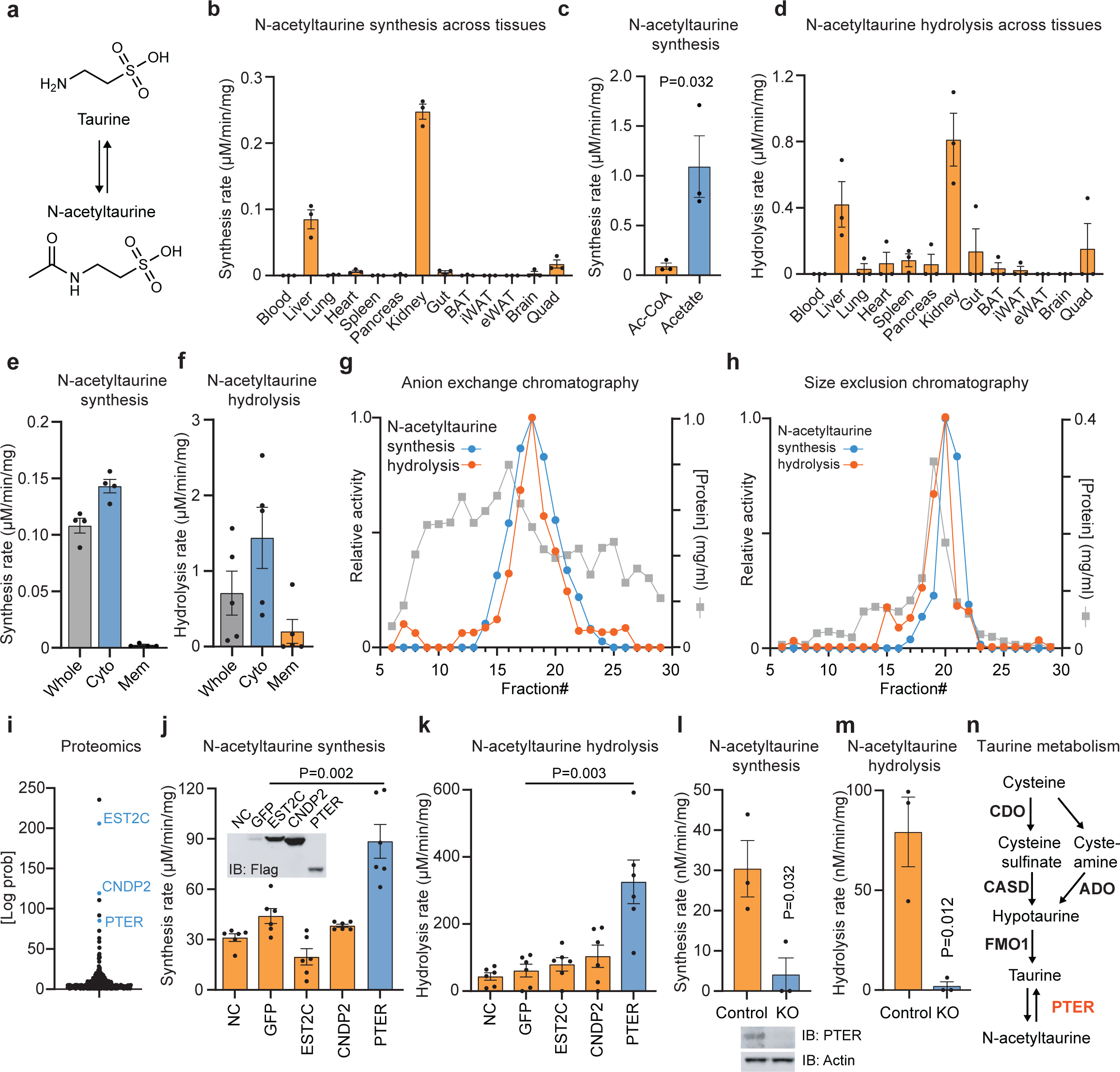
Activity-guided biochemical fractionation and proteomics identifies PTER as a taurine N-acetyltransferase/hydrolase. (a) Schematic of the biochemical interconversion of taurine and N-acetyltaurine. (b-d) Rate of N-acetyltaurine synthesis (b,c) or hydrolysis (d) activity by the indicated mouse whole tissue homogenate. Tissues were collected from 10 to 14-week-old male C57BL/6J mice (N=3/tissue). Reactions were performed using 100 µg tissue homogenates at 37°C for 1 hr with 10 mM acetate and 10 mM taurine (b), 10 mM acetyl-CoA and 10 mM taurine (c), or 100 µM N-acetyltaurine (d). (e,f) Rate of N-acetylaturine synthesis (e) and hydrolysis (f) activity in the indicated fraction of total kidney lysate. Reactions were performed as in (b). (g,h) Relative N-acetyltaurine synthesis (blue traces), hydrolysis (orange traces), and protein concentrations (grey traces) by the indicated fraction following anion exchange chromatography (g) or size exclusion chromatography (h). (i) Byonic P-values of proteins identified in fraction 20 following size exclusion chromatography. (j-m) Rate of N-acetyltaurine synthesis (j,l) or hydrolysis (k,m) activity from HEK293T cell lysates after transfection with the indicated plasmids (j,k) or from control or PTER-KO cell lysates (l,m). Reactions were conducted with 100 µg cell lysates at 37 °C for 1 hr with 10 mM acetate and 10 mM taurine (j,l) or 100 µM N-acetyltaurine (k,m). N=3/group. (j,l) insert: Western blot using an anti-Flag antibody of HEK293T cell lysates two days after the indicated transfection (j) or in WT and PTER-KO cells (l). (n) Schematic of revised taurine metabolic pathway showing the role of PTER as a bidirectional taurine N-acetyltransferase/hydrolase. For (b-f) and (j-m), data are shown as mean ± SEM. In (c), P-value were calculated by Student’s t-test. In (j-m), P-values were calculated from two-tailed unpaired t-tests. All experiments were repeated twice and similar results were obtained.

Next, we sought to purify the renal taurine N-acetyltransferase/hydrolase activity. Fractionation of kidney tissues into a cytosolic fraction and a 100,000 x g membrane fraction revealed enrichment of both N-acetyltaurine synthesis and hydrolysis activities in the cytosolic fraction (**Fig. 1e,f**). We subjected the cytosolic fraction to sequential anion exchange and size exclusion chromatography. In anion exchange, a single peak of taurine N-acetyltransferase activity could be detected that peaked in fractions #15-20; the reverse N-acetyltaurine hydrolysis activity exhibited an identical elution profile (**Fig. 1g**). The fractions with peak activity (#17-19 inclusive) were pooled and subjected to size exclusion chromatography. Again, a single peak of activity was observed that centered around fraction #20 in both synthesis and hydrolysis directions (**Fig. 1h**). These data suggest that the same renal enzyme, rather than two distinct enzymes, catalyzes both taurine N-acetyltransferase and N-acetyltaurine hydrolysis activities.

### PTER is a bidirectional taurine N-acetyltransferase/hydrolase

The active fraction #20 was analyzed by shotgun proteomics. In total, we identified 247 proteins with at least 1 peptide match (**Supplemental Table 1**). **Fig. 1i** shows the ranking of these proteins by Byonic P-values which measures the likelihood of a protein identification by random chance. The highest-ranking enzymes within this list were the esterase EST2C (CES2C, rank #2)^27^, the peptidase/synthase CNDP2 (rank #3)^28,29^, and PTER (rank #6), a putative metal-dependent hydrolase of unknown enzymatic activity or function. To directly determine whether any of these candidate enzymes could catalyze taurine N-acetylation and/or N-acetyltaurine hydrolysis in vitro, we transfected cDNAs encoding each of these three enzymes into HEK293T cells. Lysates from PTER-, but not EST2C-or CNDP2-transfected cells, exhibited higher N-acetyltaurine synthesis and hydrolysis activities compared to GFP-transfected control lysates (**Fig. 1j,k**). We also observed that GFP-transfected cells exhibited a basal taurine N-acetyltransferase/hydrolase activity over background, which we speculated might be due to the endogenous human PTER. Indeed, cell lysates from PTER-KO HEK293T cells generated via CRISPR/Cas9 exhibited complete loss of N-acetyltaurine synthesis and hydrolysis activity compared to control cells (**Fig. 1l,m**). We conclude that PTER is sufficient to confer both taurine N-acetyltransferase and N-acetyltaurine hydrolysis activities to HEK293T cell lysates and that PTER is also necessary for the endogenous taurine N-acetyltransferase/hydrolase activity in HEK293T cells. Based on these data, **Fig. 1n** shows the new biochemical assignment of PTER as a taurine N-acetyltransferase/hydrolase within the context of the endogenous biochemical pathways of taurine metabolism.

### Enzymology and mutagenesis of recombinant PTER

To determine if the entire N-acetyltransferase/hydrolase activity is encoded solely by the PTER polypeptide, or whether additional protein co-factors might be required, we next generated purified recombinant mouse PTER by heterologous expression in bacteria. Using this protein, the equilibrium constant (K) of the reversible taurine N-acetyltransferase/hydrolase reaction was 0.117 M^-1^ (**Fig. 2a**), which is comparable to equilibrium constants measured in other reversible metabolic amidase reactions^30^. Next, we performed kinetic studies with the recombinant PTER enzyme. We observed substrate concentration-dependent formation of N-acetyltaurine with Km of 11 and 64 mM and Kcat of 1000 s^-1^ and 5900 s^-1^ for acetate and taurine, respectively (**Fig. 2b,c**). In the reverse hydrolysis direction, a comparable catalytic activity (Kcat = 2600 s^-1^) was measured. The affinity between PTER and N-acetyltaurine was higher (Km = 430 μM) than that previously observed between PTER and either acetate or taurine (**Fig. 2b-d**). We conclude that the recombinant PTER polypeptide alone is sufficient to produce a functional taurine N-acetyltransferase/hydrolase enzyme without the need for additional protein co-factors. In addition, recombinant PTER exhibits kinetic characteristics similar to previously reported metabolic amidase-type enzymes.

**Fig. 2.**
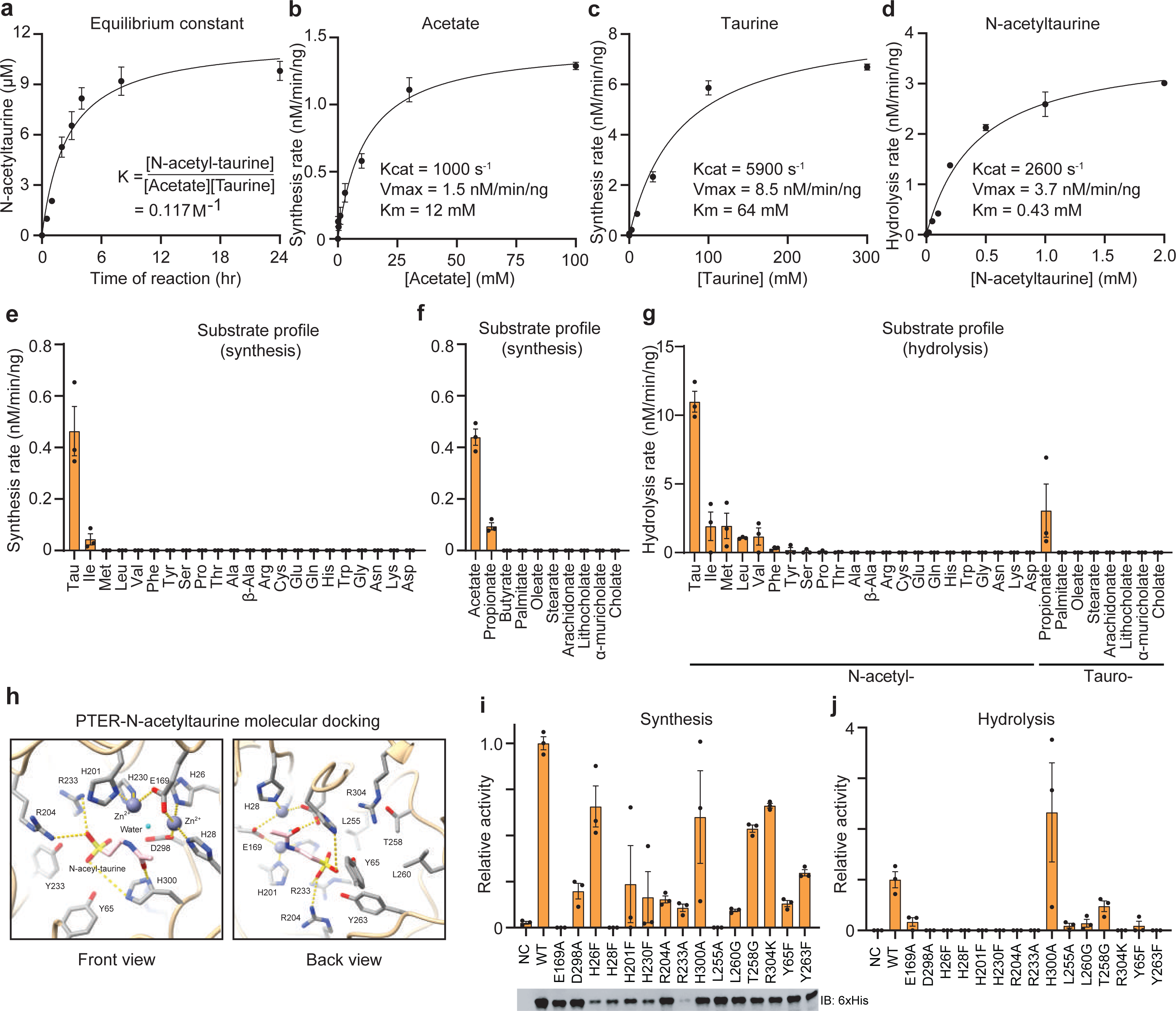
Enzymological characteristics and mutagenesis studies of recombinant mouse PTER in vitro. (a) Time-dependent formation of N-acetyltaurine following incubation of purified recombinant mPTER (100 ng) with 10 mM acetate and 10 mM taurine at 37°C. N=3/group. (b-d) Rate of N-acetyltaurine production (b,c) or hydrolysis (d) following incubation of purified recombinant mPTER with the indicated concentration of acetate and 10 mM taurine (b), the indicated concentration of taurine and 10 mM acetate (c), or the indicated concentration of N-acetyltaurine (d) at at 37°C for 1 hr. N=3/group. (e-g) Rate of synthesis (e,f) or hydrolysis (g) following incubation of purified recombinant mPTER (100 ng) with 10 mM acetate and 10 mM of indicated amino acid head group (e), 100 mM taurine and 10 or 1 mM of indicated acid (f) or 100 µM of the indicated substrate (g) at at 37°C for 1 hr. For (f), long chain fatty acids were used at 1 mM and the other organic acids were used at 10 mM. N=3/group. (h) Molecular docking of mPTER and N-acetyltaurine. Individual amino acid residues, two zinc ions (dark blue) and one water molecule (light blue) were highlighted. (I,j) Rate of N-acetyltaurine synthesis (i) or hydrolysis (j) for total bacterial lysates overexpressing the indicated mPTER mutant and Western blot using an anti-6xHIS antibody (I, bottom) of total bacterial lysates following induction of protein expression. Reactions were performed with 10 mM acetate and 10 mM taurine (i) or 100 µM N-acetyltaurine (j) at 37°C for 1 hr. N=3/group. For (a-g) and (i-j), data are shown as mean ± SEM. Data were fitted to Michaelis–Menten kinetics (solid line) using GraphPad Prism. All experiments were repeated twice and similar results were obtained.

Next, we evaluated the substrate specificity of recombinant PTER using a panel of amino acid and organic acid substrates. On the amino acid side, taurine was the best substrate when incubated with PTER and acetate (**Fig. 2e**). Little to no activity was observed when other amino acids were tested (**Fig. 2e**). On the N-acyl donor side, we also observed high PTER activity for acetate and, to a much lower extent, propionate (**Fig. 2f**). PTER did not catalyze taurine N-acylation with butyrate, longer chain fatty acids, or steroid-based acyl donors as substrates (**Fig. 2f**). Lastly, we tested the substrate specificity of PTER in the hydrolysis direction. As shown in **Fig. 2g**, PTER catalyzed robust N-acetyltaurine hydrolysis. PTER also catalyzed the hydrolysis of several other N-acetyl amino acids and N-propionyltaurine but at lower rates (<20%) compared to N-acetyltaurine. No activity was observed for most of the other substrates tested, including long chain N-fatty acyl taurines. These data demonstrate that PTER exhibits high substrate specificity for taurine, acetate, and N-acetyltaurine, but not other related acyl donors or amino acid head groups.

To determine the active site residues important for PTER enzyme activity and bidirectionality, we docked N-acetyltaurine into an Alphafold-modeled PTER^31^. For these docking and modeling purposes, we selected zinc as the divalent metal cation. In the modeled active site, we identified residues with potential interactions with N-acetyltaurine (e.g., H300, R233, R204), the metal cation (e.g, H26, H28, E169), as well as other active site residues that were proximal to the substrate (e.g, L255, Y65, T258, **Fig. 2h**). To determine the contribution of these active site side chain interactions to PTER enzyme catalysis, a total of 15 single point mutation bacterial recombinant mouse PTER proteins were produced and assayed for taurine N-acetyltransferase/hydrolase activity in vitro (**Fig. 2i,j**). In general, the expression of these point mutants, with the exception of R233A, was comparable to that of WT PTER (**Fig. 2i**). An H28F completely abolished both synthesis and hydrolysis activity; this histidine corresponds with a residue that is predicted to chelate divalent metal cations in the active site (**Fig. 2i,j**). Interestingly, we also identified several mutants, exemplified by R304K and Y263F, that maintained a moderate residual activity (∼20-60%) in the synthesis direction, but had completely abolished hydrolytic activity (**Fig. 2i,j**). Therefore, mutagenesis of specific active site residues, including H28, is sufficient to completely ablate PTER enzyme activity, while mutagenesis of others leads to functional dissociation of PTER-dependent N-acetyltaurine synthesis and hydrolysis activities.

### PTER is a physiologic N-acetyltaurine hydrolase in mice

To determine the potential physiologic relevance of PTER-dependent taurine N-acetyltransferase/hydrolase activities, we obtained global PTER-KO mice. These animals were produced by the International Mouse Phenotyping Consortium (IMPC) but had not been previously studied. Overall, PTER-KO mice were born in the expected Mendelian ratios and overtly normal in their home cage behavior. Using an anti-PTER antibody, the highest PTER protein levels were detected in liver and kidney tissues of WT mice (**Fig. 3a**), which corresponded exactly with the same tissues where we had originally detected high taurine N-acetyltransferase/hydrolase activity (**Fig. 1b,d**). As expected, complete loss of PTER protein was observed in these two tissues from PTER-KO mice (**Fig. 3a**). Similarly, kidney and liver tissues from PTER-KO mice exhibited complete loss of both taurine N-acetyltransferase and N-acetyltaurine hydrolysis activity (**Fig. 3b,c**). We conclude that PTER is the principal enzyme responsible for tissue taurine N-acetyltransferase/hydrolase activities in vivo.

**Fig. 3.**
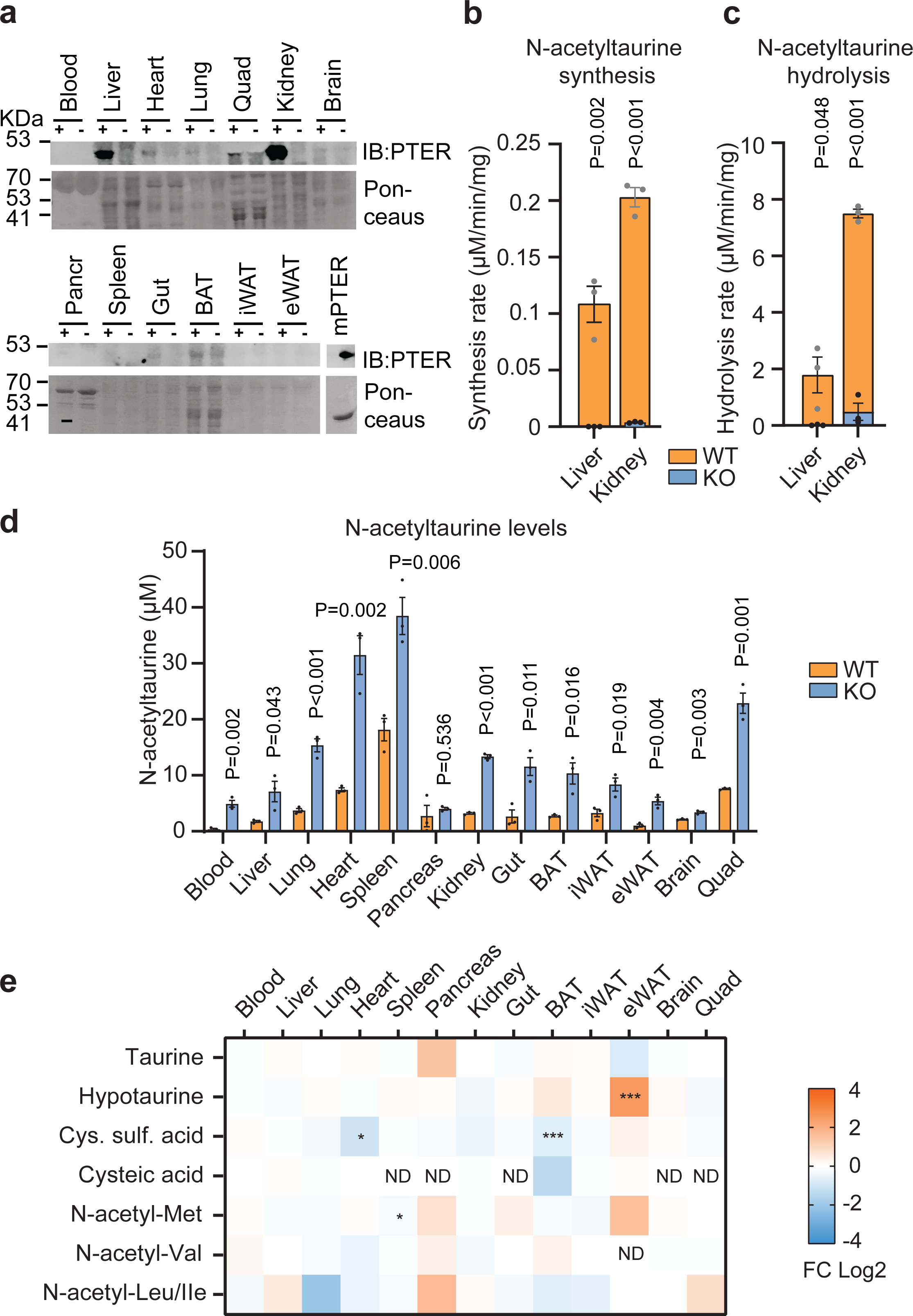
Biochemical characterization of global PTER-KO mice. (a) Anti-PTER blotting (top) and Ponceaus staining (bottom) of the indicated total tissue lysate from 4-week-old WT or PTER-KO mouse. 100 ng recombinant mPTER proteins was used as a positive control. (b,c) Rate of N-acetyltaurine production (b) or hydrolysis (c) following incubation of the indicated WT or PTER-KO total tissue lysate (100 µg) with 10 mM acetate and 10 mM taurine (b) or 100 µM N-acetyltaurine (c) at 37 °C for 1 hr. N=3/group. (d) Absolute quantitation of endogenous N-acetyltaurine levels in the indicated tissue from 4-week-old WT or PTER-KO mice. N=3/group. (e) Relative fold change (FC) of the indicated metabolites from the indicated tissue of 4-week-old WT or PTER-KO mice. N=3/group. In (b,c), data are shown as mean ± SEM. In (b-e), P-values were calculated from two-tailed unpaired t-tests. In (e), * < 0.05, ** < 0.01 and *** < 0.001.

Next, we sought to determine whether genetic PTER deficiency alters N-acetyltaurine levels in vivo and, if so, the directionality of the effect. Because other bidirectional amidases have been shown to be both physiologic synthetases as well as degradases^29,30,32^, depending on the specific substrate, it initially remained unclear whether the N-acetyltaurine would be elevated or reduced in PTER-KO mice. Using targeted metabolomics, we observed that PTER-KO tissues exhibited elevation of N-acetyltaurine which, by magnitude, ranged from 2-fold (in spleen) to >10-fold (in blood) (**Fig. 3d**). Genetic PTER deficiency therefore increases levels of N-acetyltaurine in tissues.

To understand if the elevation of N-acetyltaurine in PTER-KO mice might also result in changes to taurine levels and/or other taurine pathway metabolites, we used targeted metabolomics to measure tissue levels of taurine as well as several taurine pathway metabolites in tissues from WT and PTER-KO mice. Levels of taurine itself did not exhibit any significant genotype-dependent changes in any tissue examined (**Fig. 3e**). Hypotaurine was elevated in eWAT but not in any other tissues from PTER-KO mice; conversely, cysteine sulfinic acid was reduced in heart and brown fat mice, but not other tissues in PTER-KO mice (**Fig. 3e**). Finally, cysteic acid could be detected in a subset of tissues and its levels were unaltered in PTER-KO mice (**Fig. 3e**).

Because PTER also exhibited modest hydrolysis activity in vitro for four additional N-acetyl amino acid including N-acetylleucine, isoleucine, methionine, and valine, we used targeted metabolomics to measure the levels of these N-acetylated amino acids in WT and PTER-KO mice. As shown in **Fig. 3e**, levels of N-acetylmethionine were largely unaltered in PTER-KO tissues, except for a small reduction of N-acetylmethionine in the spleen. Levels of N-acetylvaline, N-acetylleucine and N-acetylisoleucine were also unchanged in PTER-KO mice across all tissues examined. We conclude that genetic PTER deficiency results in broad changes in N-acetyltaurine levels across all tissues, and more minor and tissue-specific changes in select taurine pathway metabolites and N-acetylmethionine.

To determine if there might be PTER-independent pathways responsible for N-acetyltaurine synthesis in mouse tissues, we measured N-acetyltaurine production in liver, kidney, brain, and blood plasma from WT and PTER-KO mice using various incubation times, buffers, and acetyl donors. Under all conditions examined, we were unable to find evidence for PTER-independent enzymatic condensation of acetate and taurine (**Extended Data Fig. S1a-d**). We were, however, able to detect a non-enzymatic condensation of acetyl-CoA with taurine (**Extended Data Fig. S1a-d**). Therefore, non-enzymatic production of N-acetyltaurine production via acetyl-CoA may be contributing to endogenous levels of N-acetyltaurine.

In urine, PTER-KO mice had ∼2-fold higher urine N-acetyltaurine compared to WT mice, with no differences in urine taurine levels (**Extended Data Fig. S1e**). N-propionyltaurine was not detectable in blood plasma (**Extended Data Fig. S1f**).

### Reduced body weight, adiposity, and food intake in PTER-KO mice

Having established PTER as the principal taurine N-acetyltransferase/hydrolase in mice, we next turned to the potential functions of this downstream biochemical pathway of taurine metabolism. Previously, Meyre et al. identified a polymorphism near the human *PTER* gene linked to early-onset and morbid adult obesity in N=14,000 European subjects^10^. Further substantiating these initial associations, in the Type 2 Diabetes Knowledge Portal the PTER gene exhibits a very strong Human Genetic Evidence (HuGE) score linked to with body mass index (BMI) (**Extended Data Fig. S2a**). These genetic data, and the prior literature of the effects of taurine supplementation on energy balance and metabolism, suggested that the PTER pathway might be involved body weight regulation.

To test this prediction, we first placed a cohort of male PTER-KO and WT littermates on high fat diet and monitored body weights and food intake over an eight-week period. After 8 weeks, food intake in PTER-KO mice was significantly reduced by a modest magnitude (∼7%) but body weight was not different (**Extended Data Fig. S2b,c**). Because taurine as a substrate for the PTER-catalyzed reaction, we reasoned that these trends in body weight and food intake in PTER-KO mice might be more robustly revealed under conditions when taurine flux is increased. We therefore placed new cohorts of WT and PTER-KO mice on a high-fat diet and also supplemented taurine in the drinking water (2.5% w/v). Under these taurine-supplemented conditions, body weights and food intake of PTER-KO mice exhibited a more marked divergence from WT mice. After 8 weeks, PTER-KO mice had lower body weight, change in body weight, and cumulative food intake compared to WT littermates (**Fig. 4a-c**). Importantly, water intake was equivalent between genotypes (**Fig. 4d**), demonstrating that the reduced food intake in PTER-KO mice was specific for nutrients rather than for all ingestion behaviors. At the end of the experiment, PTER-KO mice exhibited improved glucose tolerance and insulin sensitivity compared to WT mice, which likely represents an secondary effect to the lower body weight (**Fig. 4e,f**). Dissection of tissues revealed that the difference in body weight was due entirely to reduction of fat mass in PTER-KO mice (**Fig. 4g,h**), including lower inguinal and epididymal white adipose tissue (iWAT and eWAT, respectively), with no changes in lean mass detected. We confirmed by LC-MS that the taurine supplementation protocol increased circulating taurine levels equivalently in both WT and PTER-KO mice (**Fig. 4i**). Taurine supplementation in drinking water resulted in a hyperaccumulation of plasma N-acetyltaurine in PTER-KO mice (**Fig. 4j**).

**Fig. 4.**
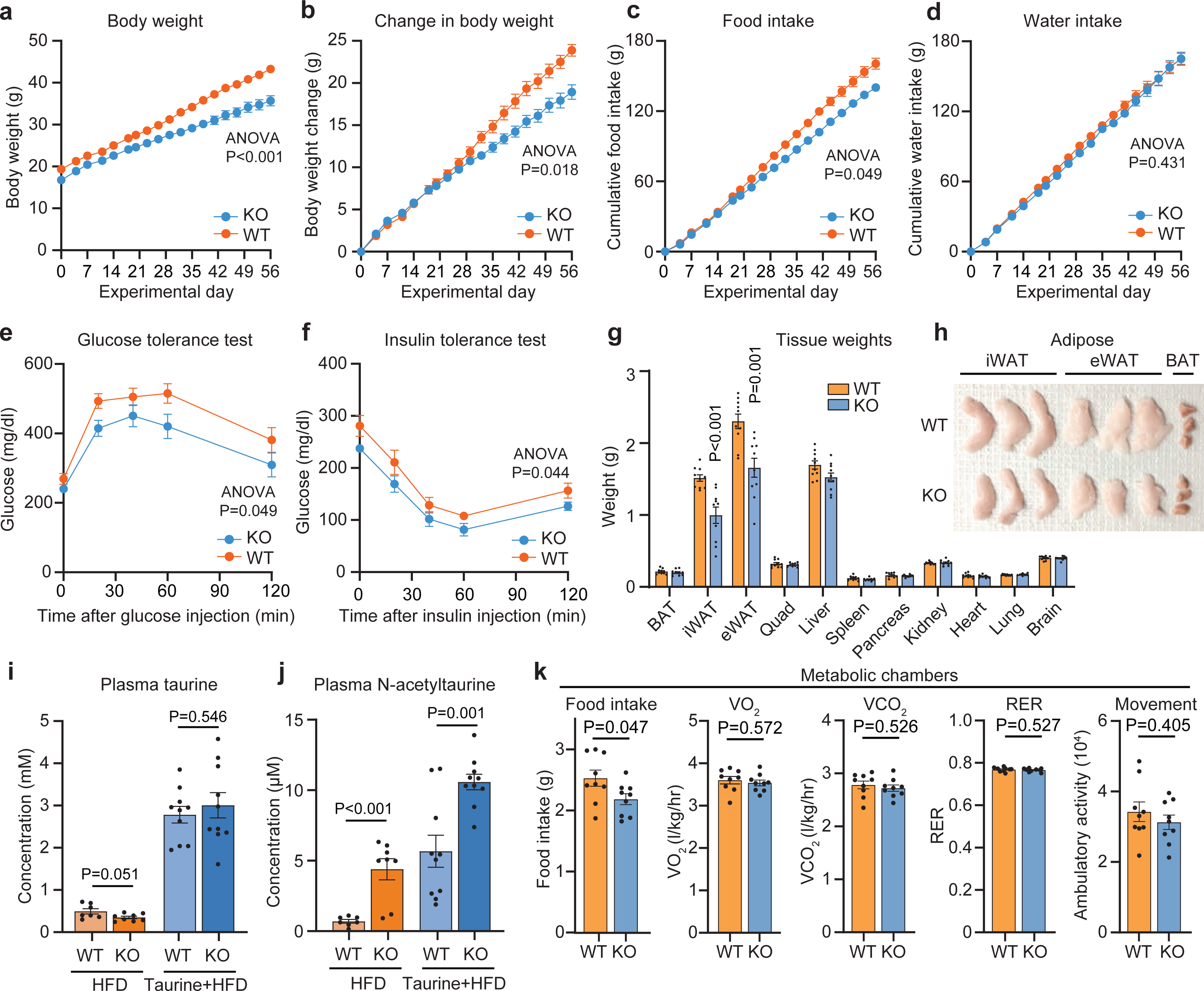
Body weight and adiposity phenotype of PTER-KO mice. (a-d) Body weight (a), change in body weight (b), cumulative food intake (c), and water intake (d) of 13 to 14-week-old male WT or PTER-KO mice on high fat diet and after taurine water supplementation (2.5% w/v). N=10/group. (e-j) Glucose tolerance test (e), insulin tolerance test (f), tissue weights (g), representative adipose tissues (h), plasma taurine levels (i), and plasma N-acetyltaurine levels (j) of 13 to 14-week-old male WT or PTER-KO mice after 8 weeks on high fat diet and taurine water supplementation (2.5% w/v). N=10/group. iWAT, inguinal white adipose tissue; eWAT, epididymal white adipose tissue; BAT, brown adipose tissue; Quad, quadriceps muscle. (k) Metabolic chamber analysis of 8 to 9-week-old-male WT or PTER-KO mice after 4 weeks of high fat diet/taurine water supplementation (2.5% w/v). N=9/group. RER, respiratory exchange ratio. Data are shown as mean ± SEM. In (a-f), P-values were calculated from two-way ANOVA with post hoc Sidak’s multiple comparisons test. In (g-k), P-values were calculated from two-tailed unpaired t-tests.

Next, we used metabolic chambers to measure parameters of whole-body energy intake and expenditure in a new cohort of PTER-KO and WT mice on taurine-supplemented water at a time point prior to the divergence in body weights (4 weeks). As expected, PTER-KO mice once exhibited reduced food intake (**Fig. 4k**). We did not observe changes in any other measured parameter including VO_2_, VCO_2_, respiratory exchange ratio or ambulatory movement (**Fig. 4k**). In an independent cohort of male WT and PTER-KO mice, metabolic chamber analysis at the end of the experiment after body weights had diverged (8 weeks) revealed reduced food intake and respiratory exchange ratio in PTER-KO mice, while VO_2_, VCO_2_, and ambulatory movement were not different between genotypes (**Extended Data Fig. S2d-i**). Non-fasted insulin levels were also not different in PTER-KO mice (**Extended Data Fig. S2j**).

Female PTER-KO on the same high fat diet/taurine-supplemented water protocol also exhibited reduced change in body weight, food intake, and adiposity compared to WT controls without any differences in water intake (**Extended Data Fig. S2k-p**). The body weight and food intake phenotype, however, was absent when either male or female PTER-KO mice were maintained on chow diet, regardless of the status of taurine-supplementation in the water (**Extended Data Fig. S3**). We conclude that PTER-KO mice have reduced adiposity, body weight, and food intake in a stimulus-dependent manner, and specifically under conditions of concurrent obesogenic diet with taurine supplementation. These data also uncover a complex gene by environment interaction of the *Pter* locus, taurine levels, and diet.

In PTER-KO and WT mice on high fat diet and taurine water, no differences in plasma GLP-1, leptin, GDF-15, ghrelin, FABP4, or adiponectin were observed at the 4-week time point (**Extended Data Fig. S4a**). At the 8-week time point, plasma leptin and plasma GDF-15 were reduced in PTER-KO mice (**Extended Data Fig. S4b**), consistent with the reduced adiposity and obesity of these animals at that time point. Protein levels of mitochondrial complexes in either liver or muscle tissues were not different between PTER-KO and WT mice at the 8-week time point (**Extended Data Fig. S4c,d**). Similarly, mRNA levels of mitochondrial or mitochondrial biogenesis markers were not different between genotypes in these two tissues (**Extended Data Fig. S4e,f**). At the 8-week time point, mRNA levels of the cytokines *Il1* and *Ccl2* were modestly reduced in adipose tissues from PTER-KO mice (**Extended Data Fig. S4g**), while mRNA levels of those same cytokines were not different in liver (**Extended Data Fig. S4h**). Liver triglycerides, AST, and ALT levels were also not different between genotypes (**Extended Data Fig. S4i,j**).

As an independent test of the stimulus-dependent body weight phenotype in PTER-KO mice, we subjected a new cohort of male WT and PTER-KO mice to a combined high fat diet and treadmill running protocol (**Extended Data Fig. S5**). We selected treadmill exercise as a second physiologic stimulus because of its previously reported effects to increase taurine levels^7,19^ (see **Methods**). We did not observe any differences in running speed or distance in WT and PTER-KO mice (**Extended Data Fig. S5a-c**). PTER-KO mice once again gained less weight and had lower food intake compared to WT mice (**Extended Data Fig. S5d-f**). Treadmill exercised PTER-KO mice also exhibited improved glucose tolerance and insulin sensitivity compared to treadmill exercised WT mice (**Extended Data Fig. S5g,h**). Dissection of tissues revealed that the weight difference was once again largely due to reductions in adipose tissue mass (**Extended Data Fig. S5i,j**). Lastly, under this the high fat diet/treadmill running protocol, we confirmed that taurine levels increased by ∼2-fold in both WT and PTER-KO mice (**Extended Data Fig. S5k**); once again, plasma N-acetyltaurine levels in the PTER-KO/exercise mice reached a level much higher than that of WT mice (with or without exercise) or even sedentary PTER-KO mice (**Extended Data Fig. S5l**).

### N-acetyltaurine administration to obese mice recapitulates the energy balance phenotype of PTER-KO mice

Because accumulation of N-acetyltaurine is the major metabolite difference between WT and PTER-KO mice, we sought to determine if N-acetyltaurine administration alone was sufficient to reproduce aspects of the energy balance phenotype in PTER-KO mice. We administered N-acetyltaurine to diet-induced obese (DIO) mice (1-50 mg/kg/day, intraperitoneally). After a single administration of N-acetyltaurine, we observed robust increases in plasma N-acetyltaurine levels that peaked at a concentration of ∼30 μM (at the 15 mg/kg dose) and ∼60 μM (at the 50 mg/kg dose) one hour after dosing (**Extended Data Fig. S6a**) without any changes to plasma taurine levels (**Extended Data Fig. S6b**). Upon chronic daily dosing, DIO mice treated with N-acetyltaurine exhibited dose-dependent reduction of both body weight (**Fig. 5a**) and food intake (**Fig. 5b**). In lean mice, N-acetyltaurine also suppressed food intake and body weight, but with a magnitude that was more moderate compared to the effect observed in DIO mice (**Extended Data Fig. S6c,d**). To determine if the effect of N-acetyltaurine required the intact amidated conjugate, we performed head-to-head comparisons of the effects of N-acetyltaurine with either acetate alone or taurine alone all at the same dose (15 mg/kg/day). Once again, N-acetyltaurine-treated mice exhibited reduced food intake and body weight, whereas mice treated with either acetate alone or taurine alone were indistinguishable from vehicle-treated mice (**Fig. 5c,d**). We conclude that administration of N-acetyltaurine to wild-type, DIO mice is sufficient to reduce body weight and food intake.

**Fig. 5.**
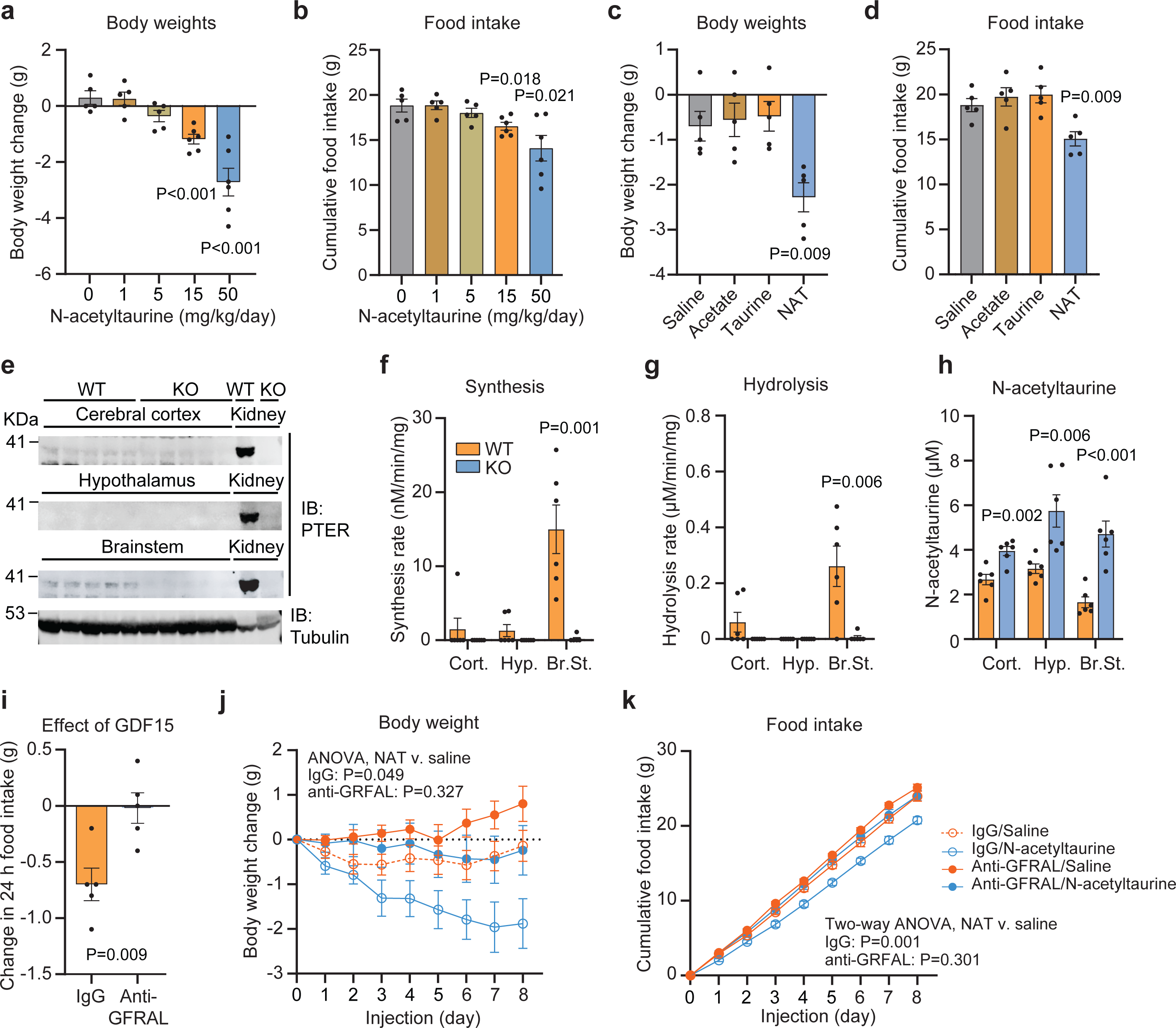
Effect of N-acetyltaurine administration to diet-induced obese mice. (a,b) Change in body weight (a) and cumulative food intake (b) of 26 to 28-week-old male DIO C57BL/6J mice following 7 days of treatment with the indicated dose of N-acetyltaurine (intraperitoneal injection). N=5/group for vehicle, 1, and 5 mg/kg/day; N=6/group for 15 and 50 mg/kg/day. (c,d) Change in body weight (c) and cumulative food intake (d) of 19 to 21-week-old male DIO C57BL/6J mice following treatment with the indicated metabolite at a dose of 15 mg/kg/day (IP) N=5 per group. NAT, N-acetyltaurine. (e-h) Western blotting with anti-PTER (top) and anti-tubulin (bottom) antibodies (e), N-acetyltaurine synthesis activity (f), N-acetyltaurine hydrolysis activity (g), and tissue N-acetyltaurine levels (h) from cortext (Cort.), hypothalamus (Hyp.), or brainstem (Br.St.) of WT or PTER-KO mice. N=4/group for (e) and N=6/group for (f-h). (i) Change in 24 h food intake of 6-month-old male DIO mice treated with a single dose of GDF15 (0.1 mg/kg, IP) in the presence of anti-GFRAL antibody (10 mg/kg, IP) or IgG control antibody (10 mg/kg, IP). N=5/group. (j,k) Change in body weight (j) and cumulative food intake (k) of 16-week old male DIO mice following saline or N-acetyltaurine (15 mg/kg/day, IP) treatment and with IgG or anti-GFRAL antibody co-treatment (10 mg/kg, IP, once every 3 days). N=10/group. Data are shown as mean ± SEM. In (a-d) and (f-i), P-values were calculated from two-tailed unpaired t-tests. In (j,k), P-values were calculated from two-way ANOVA with post hoc Sidak’s multiple comparisons test.

To better understand how N-acetyltaurine controls feeding behaviors, we examined the expression of PTER protein in various brain regions by Western blotting using our anti-PTER antibody. PTER protein was detected in the brainstem, but not hypothalamus or cerebral cortex (**Fig. 5e**). We also detected a PTER-dependent N-acetyltaurine synthesis/hydrolysis activity and accumulation of N-acetyltaurine in the brainstem. (**Fig. 5f,g**). While N-acetyltaurine was elevated in all brain regions examined, the greatest fold change was observed in the brainstem (**Fig. 5h**). Profiling of mRNA for neuropeptide and feeding related genes in both brainstem and hypothalamus did not reveal any obvious PTER-dependent changes of large magnitude (**Extended Data Fig. S7**).

Because of the established role of brainstem-restricted GDF15/GFRAL signaling in the feeding control, we tested whether the anorexigenic and anti-obesity effects of N-acetyltaurine administration requires an intact GFRAL receptor. We obtained a neutralizing anti-GFRAL antibody (IgG clone 8A2, Eli Lilly & Co.) and an IgG control antibody. We confirmed that anti-GFRAL antibody completely abrogated the anorexigenic effect of recombinant GDF15 (**Fig 5i**). As expected, N-acetyltaurine lowered body weight and food intake when co-administered with the IgG control antibody (**Fig. 5j,k**). By contrast, N-acetyltaurine did not significantly reduce either body weight or food intake in the presence of anti-GFRAL antibody (**Fig. 5j,k**). We also tested the role of GLP-1R and hypothalamic MC4R signaling in the anti-obesity effects of N-acetyltaurine. The GLP-1R antagonist Exendin-3 blocked the effects of GLP-1 peptide in food intake and body weight; however, under these conditions Exendin-3 did not blunt the body weight-lowering effect of N-acetyltaurine (**Extended Data Fig. S8a-d**). Similarly, N-acetyltaurine also suppressed food intake and body weight in MC4R-KO mice (**Extended Data Fig. S8e,f**). We conclude that PTER is expressed in the brainstem and that the full anorexigenic and anti-obesity effects of N-acetyltaurine require functional GFRAL receptors.

To determine the direct versus indirect effects of N-acetyltaurine in adipose tissues, we examined the effects of N-acetyltaurine in isolated adipocytes in vitro and after administration to mice in vivo. In vitro, N-acetyltaurine did not acutely stimulate adipocyte lipolysis as measured by glycerol release (**Extended Data Fig. S9a**). N-acetyltaurine also did not stimulate the expression of lipogenesis or lipid uptake-associated genes in isolated adipocytes (**Extended Data Fig. S9b**). A single administration of N-acetyltaurine to mice did not stimulate lipolysis or alter lipogenesis or lipid uptake gene expression in epididymal fat tissues (**Extended Data Fig. S9c,d**). Therefore N-acetyltaurine does not directly regulate lipid metabolism in adipocytes. In plasma from PTER-KO mice, we did not observe any changes in specific plasma free fatty acid species, while plasma glycerol levels were modestly increased (**Extended Data Fig. S9e-i**). In epididymal fat from PTER-KO mice, complex bidirectional changes in mRNA levels for lipid uptake and lipogenesis genes was observed, and p-HSL was slightly reduced (**Extended Data Fig. S9j,k**), all of which likely represent secondary effects due to reduced food intake in N-acetyltaurine-treated mice.

### The gut microbiome contributes to circulating N-acetyltaurine levels

Lastly, we considered the possibility that the gut microbiome may also be involved in host N-acetyltaurine metabolism. Indeed, wild-type mice treated with an antibiotics cocktail for one week exhibited a ∼30% reduction in circulating N-acetyltaurine levels without any changes in circulating taurine levels (**Fig. 6a-c**). Conversely, plasma N-acetyltaurine, but not taurine, was increased by ∼80% after colonization of germ-free mice with the defined microbial community hCom2 (**Fig. 6d,e**)^33^. Using the biochemical assay for a taurine N-acetyltransferase activity, we detected robust production and secretion of N-acetyltaurine by the cellular fraction of feces isolated from hCOM2-colonized, but not germ-free mice (**Fig. 6f**).

**Fig. 6.**
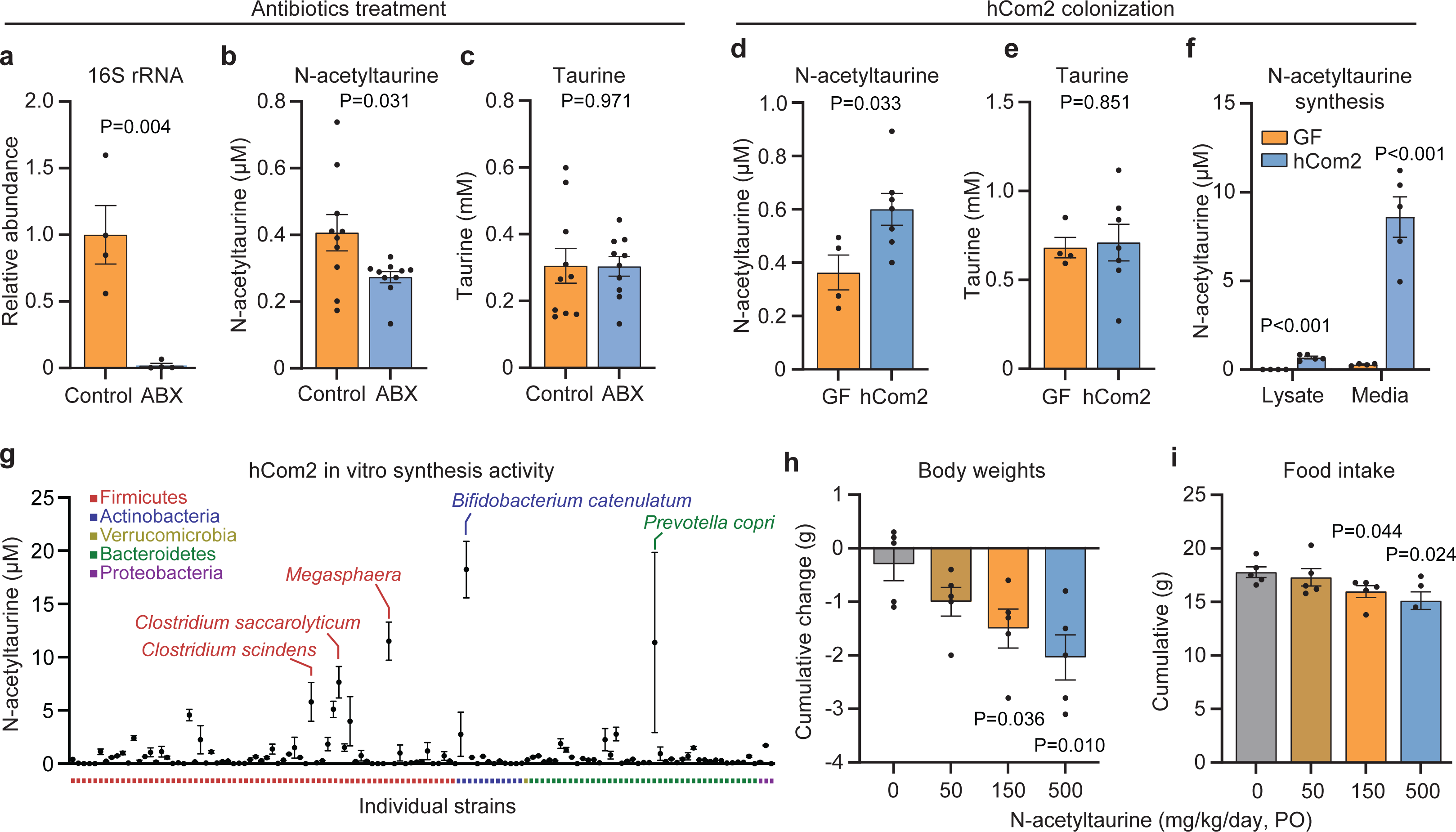
Gut microbiome contributions to circulating N-acetyltaurine levels. (a-c) Fecal 16S rRNA (a), plasma N-acetyltaurine (b), and plasma taurine (c) levels of control and antibiotic (ABX)-treated 12-to 14-week-old wild-type lean male mice. Antibiotic mixture ( chloramphenicol, spectinomycin dihydrochloride pentahydrate, apramycin sulfate, tetracycline hydrochloride, kanamycin and ampicillin at 1g/l per antibotic) was administered in drinking water ad libitum and orally gavaged (0.5 ml) every other day for a whole duration of 2 weeks. In (a), N=4/group. In (b,c), N=10/group. (d,e) Plasma N-acetyltaurine (d) and plasma taurine (e) levels of germ-free (GF) or hCom2-colonized 12-to 14-week-old wild-type lean male mice. N=4 for GF mice and N=4 for hCom2-colonized mice. (f) N-acetyltaurine levels in cell lysates or media of cells isolated from feces of GF or hCom2-colonized 12-to 14-week-old wild-type lean male mice. N=4 for GF mice and N=5 for hCom2-colonzied mice. (g) Media N-acetyltaurine levels following incubation of taurine (10 mM) and acetate (10 mM) for 48 h with individual bacterial strains from hCom2. N=2/strain. (h,i) Change in body weight (h) and cumulative food intake of 3-month-old male DIO mice treated orally with the indicated dose of N-acetyltaurine (50-500 mg/kg/day, PO). Data are shown as mean ± SEM. In (a-f) and (h,i), P-values were calculated from two-tailed unpaired t-tests.

To determine the identity of bacteria that contribute to N-acetyltaurine synthesis, we assayed individual strains of hCom2 (**Supplemental Table 2**) for production of N-acetylaturine after overnight incubation with taurine and acetate. The highest producer identified in this biochemical assay was *Bifidobacterium catenulatum* (**Fig. 6g**). Several firmicutes (*Megasphaera, Clostridium saccarolyticum,* and *Clostridium scindens*) as well as the bacteroidetes strain *Prevotella copri*, also exhibited N-acetyltaurine synthesis activity. Therefore multiple bacterial strains can contribute to N-acetyltaurine production.

To determine if N-acetyltaurine could cross the intestinal barrier intact, we administered N-acetyltaurine by oral gavage (50-500 mg/kg, PO). Under these conditions, plasma N-acetyltaurine levels dose-dependently increased to levels comparable to that found after intraperitoneal administration (**Extended Data Fig. S10**). In addition, oral administration of N-acetyltaurine to DIO mice (50-500 mg/kg/day, PO) dose-dependently reduced both food intake and body weight (**Fig. 6h,i**). We conclude that the gut microbiome is a source of circulating N-acetyltaurine in the host. In addition, oral administration of N-acetyltaurine is sufficient to reduce food intake and body weight.

## Discussion

Here we provide multiple independent lines of evidence that show PTER is a taurine N-acetyltransferase/hydrolase that controls food intake and body weight under physiologic or dietary stimuli that increase taurine levels. First, by activity-guided fractionation, PTER is detected in a partially purified fraction from mouse kidney with taurine N-acetyltaurine/hydrolase activity. Second, recombinant PTER is a bidirectional mammalian taurine N-acetyltransferase/hydrolase in vitro. Third, genetic ablation of PTER in mice results in complete loss of taurine N-acetyltransferase/hydrolase activities and concomitant elevation of N-acetyltaurine across multiple tissues. PTER-KO mice also exhibit lower body weight and adiposity upon nutritional or physiologic stimuli that increase taurine levels. Lastly, N-acetyltaurine is sufficient to suppress food intake, adiposity, and body weight in diet-induced obese mice in a manner that depends on functional GFRAL receptors.

Until now, PTER has been an enigmatic and poorly studied enzyme that was largely uncharacterized with respect to both biochemical activity and physiologic function. Bacterial phosphotriesterases (PTEs) catalyze the hydrolysis of organophosphate substrates; however, the biochemical activity of mammalian PTER, and their endogenous substrates in vivo, were unknown. In addition, genetic associations linked the human PTER locus with body mass index, but the causal and mechanistic basis underlying this genetic association was also unknown. Our data provide a coherent and unifying model that answers these questions. First, we show that PTER is a central and previously unrecognized node in taurine and acetate metabolism. To the best of our knowledge, PTER is the first enzyme reported to catalyze taurine N-acetylation and hydrolysis. Second, we demonstrate that genetic loss of PTER, or pharmacological administration of N-acetyltaurine, results in reduced food intake, adiposity, and body weight. These data show that PTER is causally linked to body weight and establish a role for the PTER-regulated metabolite N-acetyltaurine in this process.

In addition to regulation of steady state N-acetyltaurine levels by PTER, we also provide evidence that the gut microbiome can modulate plasma levels of N-acetyltaurine. In the future, additional pathways involved in the biosynthesis and/or metabolism of N-acetyltaurine might be revealed by a more careful survey of in vitro enzyme assay conditions beyond those tested here. In addition, turnover flux measurements of taurine and N-acetyltaurine, especially under diverse physiologic stimuli, and in WT or PTER-KO mice, would be valuable to understanding the kinetics and dynamic regulation of these metabolites in vivo. That N-acetyltaurine is under co-regulation by both host as well as microbial pathways also raises the possibility that rational manipulation of the gut microbiome may also be a viable strategy for augmenting host N-acetyltaurine levels to reduce body weight.

A major unanswered question is the precise molecular and circuit mechanisms by which N-acetyltaurine regulates feeding behaviors and energy balance. Our data point to the brainstem, and GFRAL receptors in particular, as important downstream effectors of N-acetyltaurine. However, N-acetyltaurine likely does not directly bind GFRAL itself because N-acetyltaurine is a metabolite and consequently does not share any structural similarity with the natural GFRAL ligand GDF15. The specific pathway of the crosstalk between the PTER/N-acetyltaurine and GFRAL pathways may be complex and involve intermediate steps. For example, N-acetyltaurine may modulate neurotransmission pathways directly since this metabolite shares structural similarity with the neurotransmitter acetylcholine and taurine itself has been shown to be an agonist of GABAA and glycine receptors^2^. Alternatively, N-acetyltaurine may indirectly affect neurotransmission pathways by metabolically altering and/or diverting acetate units away from the pool of acetylated signaling molecules (e.g., acetylcholine or melatonin). The precise downstream pathways of N-acetyltaurine anorexigenic action remains an important area of future work. Conditional *Pter* alleles, which are currently being developed in our laboratory, will enable dissection of the central versus peripheral contributions of PTER to the whole-body energy balance phenotypes. In addition, future generation of knock-in mice with synthesis-only or hydrolysis-only PTER mutations would enable functional dissection of these two enzymatic activities in vivo.

In recent years there has been an explosion of interest in taurine and taurine supplementation for many other aspects of human health and disease beyond metabolism. For instance, taurine has been recently linked to multiple age-associated phenotypes.^19,34,35^ Our data show that that taurine-derived metabolites are not simply biomarkers or inert byproducts, but in fact chemical effectors of the elevated taurine state. Future studies exploring the role of taurine metabolites such as N-acetyltaurine in these other processes may identify opportunities where pharmacological manipulation of secondary taurine metabolism may be therapeutically useful.

## Methods

### Chemicals

DL-Dithiothreitol (DTT) (D0632-1G), taurine (T0625-100G), acetate (S2889-250G), isoleucine (I2752-1G), L-methionine (M5308-25G), L-leucine (L8000-25G), L-valine (V-0500), L-serine (S260-0), L-proline (P0380-100G), L-threonine (T8625-1G), L-alanine (A7627-1G), β-alanine (05160-50G), L-arginine (A5006-100G), L-cysteine (168149-25G), L-glutamic acid (49621-250G), L-glutamine (G-3126), L-histidine (H-8000), L-tryptophan (T0254-5G), L-asparagine (A0884-25G), L-lysine (L5501-5G), acetate (S2889-250G), propionate (P1880-100G), butyrate (B5887-1G), palmitate (P9767-5G), oleate (O7501-1G), stearate (S3381-5G), arachidonate (10931), N-acetyl-L-methionine (01310-5G), N-acetyl-L-leucine (441511-25G), N-acetyl-L-phenylalanine (857459-5G), N-acetyl-L-tyrosine (PHR1173-1G), N-acetyl-L-serine (A2638-1G), N-acetyl-L-proline (A0783-1G), N-acetyl-L-alanine (A4625-1G), N-acetyl-L-arginine (A3133-5G), N-acetyl-L-cysteine (A7250-25G), N-acetyl-L-glutamic acid (855642-25G), N-acetyl-L-glutamine (A9125-25G), N-acetyl-L-tryptophan (A6376-10G), N-acetyl-glycine (A16300-5G), N-acetyl-L-asparagine (441554-1G), N-acetyl-L-lysine (A2010-1G), N-acetyl-L-aspartic acid (00920-5G), chloramphenicol (C0378-25G), spectinomycin dihydrochloride pentahydrate (S4014-25G), apramycin sulfate (A2024-5G), tetracycline hydrochloride (T7660-5G), and ampicillin (A9518) were purchased from Sigma. Paraformaldehyde (AAJ19943K2), tryptone (BP1421-500), yeast extract (BP1422-500), L-tyrosine (A11141.22), glycine (G48-212), N-acetyl-β-alanine (H50208.03), and kanamycin (11815032) were purchased from ThermoScientific. N-acetyltaurine (35169), lithocholate (20253), α-muricholate (20291) and taurocholate (16215), N-palmitoyl-taurine (10005611), N-oleoyl-taurine (10005609), N-stearoyl-taurine (10005610), N-arachidonoyl-taurine (10005537), taurolithocholic acid (17275), tauro-α-muricholic acid (20288) and taurocholic acid (16215) were purchased from Cayman. L-phenylalanine were purchased (A13238), N-acetyl-L-isoleucine (H66771), N-acetyl-L-valine (H66943) and N-acetyl-L-histidine (Alfa Aesar, J65657) were purchased from Alfa Aesar. L-aspartic acid (11625) was purchased from United States Biochemical Corporation. N-acetyl-L-threonine (03262) was purchased from CHEM-IMPEX INT’L INC, Heavy N-acetyltaurine and N-propionyl-taurine were synthesized by Acme. GLP-1 (7-37) peptides (CP0005) were purchased from Genescript. Exendin-3 (9-39) amide (2081) was purchased from Tocris. Recombinant GDF15 (957-GD) was purchased from R&D systems. Anti-GFRAL neutralizing antibody and control IgG antibody^36^ were obtained from Eli Lilly, a gift generously provided by Dr. Emmerson.

### Cell line cultures

HEK293T cell line was obtained from the American Type Culture Collection (ATCC) and grown at 37 °C with 5% CO2. The culture medium consists of Dulbecco’s modified Eagle’s medium (Corning, 10-017-CV) with 10% FBS (Corning, 35010CV) and 1:1000 penicillin–streptomycin (Gibco, 15140-122). For transient transfection, cells were transfected in 10 cm^2^ at ∼60% confluency using PolyFect (Qiagen, 301107) and washed with complete culture medium 6 h later. The HEK293T cells were negative following testing for mycoplasma contamination.

### Generation of PTER-KO cells

The pLentiCRISPRv2 system was used to generate PTER-KO HEK293T cells. The single guide RNA (sgRNA) used was 5′-GATGGAACCAGTATCAAGTG-3′. The following oligonucleotides were used to clone the sgRNA into the plentiCRISPRv2 vector: forward, 5′-CACCGGATGGAACCAGTATCAAGTG-3′; reverse, 5′-AAACCACTTGATACTGGTTCCATCC-3′.

Lentiviral particles were produced in the HEK293T cell line using Polyfect for the co-transfection of the cloned plentiCRISPRv2 plasmid with the viral packing psPAX2 plasmid and the viral envelope pMD2.G plasmid. A plentiCRISPRv2 plasmid without any sgRNA insert was used as a negative control. Medium containing lentivirus was collected 48 h after transfection and filtered through a 0.45-µM filter. The supernatant was then mixed in a 1:1 ratio with polybrene (Sigma, TR-1003-G) to a final concentration of 8 µg/ml polybrene. The viral mixture was added to HEK293T cells at 40–50% confluence in 6-well plates. Transduced cells were transferred to a 10 cm² plate and subjected to puromycin selection for a period of 3-6 days. Surviving cells were then trypsinized, resuspended and plated at a 10,000x dilution to a new 10 cm^2^ plate. Two weeks later, individually distinguishable colonies were visually identified and then transferred to a 96-well plate using a sterile pipette tip. Finally, single HEK293T cell clones exhibiting complete loss of endogenous PTER protein were confirmed via Western blotting using a polyclonal anti-PTER antibody (Invitrogen, TR-1003-G).

### Western blotting

For analyzing samples from cell culture, cells were collected and lysed by probe sonication. Cell lysates were centrifuged at 13,000 rpm for 10 min at 4 °C. The supernatant was collected, boiled for 10 min at 95 °C in 4× NuPAGE LDS Sample Buffer (ThermoFisher, NP0008) supplemented with 100 mM DTT (Sigma, D0632-1G). For analyzing samples of mice, blood was obtained through submandibular bleeding using a 21G needle (BD, 305129) into lithium heparin tubes (BD, 365985). Blood was subsequently spun down at 5,000 rpm for 5 min at 4 °C to retrieve the supernatant plasma fractions. All tissues were dissected, weighed on a scale, collected into Eppendorf tubes, and immediately frozen on dry ice and stored at −80 °C. A stereotaxic device was used to dissect out hypothalamus and brainstem. Adipose tissues were preserved in 4% paraformaldehyde (FisherScientific, AAJ19943K2) for histology analysis. Tissues were then mixed with 0.5 ml of cold RIPA buffer and homogenized using a Benchmark BeadBlaster Homogenizer at 4 °C. The mixture was spun down at 13,000 rpm for 10 min at 4 °C to pellet the insoluble materials. The supernatant was quantified using a tabletop Nanodrop One or using a BCA Protein Assay Kit (FisherScientific, 23250) and analyzed by western blot. Adipose tissues from DIO mice were separately processed using a protein extraction kit to remove lipids (Invent Biotechnologies, AT-022). Proteins were separated on NuPAGE 4–12% Bis-Tris gels and transferred to nitrocellulose membranes. Equal loading was ensured by staining blots with Ponceau S solution. Blots were then incubated with Odyssey blocking buffer for 30 min at room temperature and incubated with primary antibodies (1:1000 dilution rabbit anti-PTER antibody (Invitrogen, PA5-20750), 1:5000 dilution rabbit anti-β-actin antibody (Abcam, ab8227), 1:1000 dilution mouse anti-OxPhoS cocktail antibody (Invitrogen, 45-8099), 1:1000 dilution rabbit anti-HSL antibody (Novus biologicals, NB110-37253), 1:1000 dilution rabbit anti-pHSL (Novus biologicals, NBP3-05457), 1:1000 dilution rabbit anti-ATGL (Cell signaling, 2138), 1:1000 1:5000 dilution mouse anti-α-tubulin antibody (Cell Signaling, 3873S), 1:5000 dilution mouse anti-Flag antibody (Sigma, F1804-200UG), 1:1000 dilution rabbit anti-6xHis antibody (Abcam, ab9108)) in blocking buffer overnight at 4 °C. Blots were washed three times with PBST (0.05% Tween-20 in PBS) and stained with species-matched secondary antibodies (1:10000 dilution goat anti-rabbit IRDye 800RD (LI-COR, 925-68070) and 1:10000 dilution goat anti-mouse IRDye 680RD (LI-COR, 925-68070)) at room temperature for 1 h. Blots were further washed three times with PBST and imaged with the Odyssey CLx Imaging System.

### Generation of recombinant mPTER proteins

mPTER gene (Uniprot Q60866) was codon optimized to ensure bacterial expression and was synthesized as gBlocks with IDT. The gene fragment was then inserted into the pET-20b vector containing a C-terminal hexa-Histidine (His) tag. DNA sequences encoding a Strep tag were cloned into the N-terminus of mPTER for Strep-Tactin-based purification. BL21 competent bacteria (ThermoScientific, EC0114) were used to transform pET-20b-mPTER plasmids and subsequently cultured in LB medium with ampicillin at 37 °C on a shaker overnight. BL21 cells were then transferred to autoinduction medium, which consisted of the following components: 10 g tryptone (FisherScientific, BP1421-500), 5 g yeast extract (FisherScientific, BP1422-500), 2 ml MgSO_4_ (1 M), 1 ml metal solution (0.05 M Feccir citrate, 0.02 M CaCl_2_, 0.02 M ZnSO_4_, 2 µM CoCl_2_, 2 µM CuSO_4_, 2 µM NiCl_2_, 2 µM Na_2_MoO_4_, 2 µM Boric acid), 20 ml salt solution (167.5g Na_2_HPO_4_, 85g KH_2_PO_4_, 53.4g NH_4_Cl and 17.8g Na_2_SO_4_ in 500 ml water in total) and 20 ml sugar solution (125 g glycerol, 12.5 g glucose and 50 g α-lactose in 500 ml water in total) in a total volume of 1 L. The bacteria were cultured until the optical density value reached a range of 0.5 to 0.7. Bacteria were subsequently incubated at 15 °C overnight before being spun down at 8,000 rpm for 30 min at 4 °C. Bacteria were then lysed in PBS through probe sonication on ice to release cytosolic proteins. Soluble fractions were isolated via high-speed centrifugation at 15,000 rpm for 30 min at 4 °C. And they were run down a Nickel column using an ӒKTA pure™ chromatography system. The elution was performed from 0 mM to 300 mM NaCl in PBS over a gradient involving 60 column volumes. Fractions containing mPTER proteins were pooled together before undergoing another round of purification. This step involved running fractions down columns loaded with Strep-Tactin resins (IBA, 2-1208-002), following the manufacturer’s instructions. The bound mPTER proteins were eluted by 2.5 mM D-Desthiobiotin before passing through a HiPrep 16/60 Sephacryl S-200 size-exclusion column (Sigma, GE17-1166-01) in buffer containing 25 mM Tris and 100 mM NaCl. Finally fractions containing monomeric mPTER recombinant proteins were pooled together and subjected to SDS-PAGE gel electrophoresis to ensure >95% purity was achieved. The recombinant proteins were aliquoted and stored at −80 °C for subsequent enzymatic assays.

### Enzymatic assays

A total of 100 μg of proteins derived from cell or tissue lysates, or 100 ng of recombinant mPTER proteins, or 50 µl of chromatography fractions were subjected to incubation in a 50 µl PBS solution at 37°C for 1 hour. For assays using kidney membrane and soluble fractions, total kidney homogenates were transferred into ultracentrifuge inserts and spun at 100,000 x g on a Beckman Centrifuge I8-70M for 1 hr at 4°C. The supernatant was quantified as the kidney soluble fraction and the pellet was resuspended thoroughly in PBS and measured using a using a tabletop Nanodrop One. 10 mM taurine (Sigma, T0625-100G) and 10 mM acetate (Sigma, S2889-250G) were added for N-acetyltaurine synthesis. 100 µM N-acetyltaurine (Cayman, 35169) was added for assaying hydrolysis. For assays testing the substrate scope of mPTER synthesis, 10 mM L-isoleucine (Sigma, I2752-1G), L-methionine (Sigma, M5308-25G), L-leucine (Sigma, L8000-25G), L-valine (Sigma, V-0500), L-phenylalanine (Alfa Aesar, A13238), L-tyrosine (ThermoScientific, A11141.22), L-serine (Aldrich Chemical Company Inc, S260-0), L-proline (Sigma, P0380-100G), L-threonine (Sigma, T8625-1G), L-alanine (Sigma, A7627-1G), β-alanine (Sigma, 05160-50G), L-arginine (Sigma, A5006-100G), L-cysteine (Sigma, 168149-25G), L-glutamic acid (Sigma, 49621-250G), L-glutamine (Sigma, G-3126), L-histidine (Sigma, H-8000), L-tryptophan (Sigma, T0254-5G), glycine (FisherChemical, G48-212), L-asparagine (Sigma, A0884-25G), L-lysine (Sigma, L5501-5G), L-aspartic acid (United States Biochemical Corporation, 11625) were individually incubated with 10 mM acetate (Sigma, S2889-250G); 10 mM propionate (Sigma, P1880-100G), 10 mM butyrate (Sigma, B5887-1G) was incubated with 10 mM taurine; 1 mM palmitate (Sigma, P9767-5G), oleate (Sigma, O7501-1G), stearate (Sigma, S3381-5G), arachidonate (Sigma, 10931), lithocholate (Cayman, 20253), α-muricholate (Cayman, 20291) and taurocholate (Cayman, 16215) were individually incubated with 100 mM taurine. For assays testing the substrate scope of mPTER hydrolysis, 100 μM N-acetyl-L-isoleucine (Alfa Aesar, H66771), N-acetyl-L-methionine (Sigma, 01310-5G), N-acetyl-L-leucine (Sigma, 441511-25G), N-acetyl-L-valine (Alfa Aesar, H66943), N-acetyl-L-phenylalanine (Sigma, 857459-5G), N-acetyl-L-tyrosine (Sigma, PHR1173-1G), N-acetyl-L-serine (Sigma, A2638-1G), N-acetyl-L-proline (Sigma, A0783-1G), N-acetyl-L-threonine (CHEM-IMPEX INT’L INC, 03262), N-acetyl-L-alanine (Sigma, A4625-1G), N-acetyl-β-alanine (ThermoScientific, H50208.03), N-acetyl-L-arginine (Sigma, A3133-5G), N-acetyl-L-cysteine (Sigma, A7250-25G), N-acetyl-L-glutamic acid (Sigma, 855642-25G), N-acetyl-L-glutamine (Sigma, A9125-25G), N-acetyl-L-histidine (Alfa Aesar, J65657), N-acetyl-L-tryptophan (Sigma, A6376-10G), N-acetyl-glycine (Sigma, A16300-5G), N-acetyl-L-asparagine (Sigma, 441554-1G), N-acetyl-L-lysine (Sigma, A2010-1G), N-acetyl-L-aspartic acid (Sigma, 00920-5G), N-propionyl-taurine (Acme, AB38328), N-palmitoyl-taurine (Cayman, 10005611), N-oleoyl-taurine (Cayman, 10005609), N-stearoyl-taurine (Cayman, 10005610), N-arachidonoyl-taurine (Cayman, 10005537), taurolithocholic acid (Cayman, 17275), tauro-α-muricholic acid (Cayman, 20288) and taurocholic acid (Cayman, 16215) were used. Reactions were then quenched and metabolites were extracted by 150 µl of a 2:1 mixture of acetonitrile:methanol. The mixture was spun down at 15,000 rpm for 30 min at 4 °C. The supernatant was subsequently transferred to mass spec vials and ready for LC-MS analysis.

### Molecular docking

The AlphaFold-predicted structure of murine PTER (AF-Q60866-F1) was used to search for proteins with structural or sequence homology, using FoldSeek and Blast respectively. The top-predicted structural match from the PDB as identified by FoldSeek was PDB 3K2G, a Resiniferatoxin-binding protein isolated from Rhodobacter sphaeroides. This crystal structure, along with annotation in uniprot, and metal binding-site prediction using MIB2, all indicated the presence of 2 zinc ions in the active site of PTER. Molecular docking was performed with CB-Dock2, an online docking server using curvature-based cavity prediction followed by AutoDock Vina-based molecular docking. The substrate compounds N-acetyltaurine was prepared as a SDF file, and the AlphaFold-predicted protein structure for PTER was prepared as a PDB file. Ligand-receptor docking was performed using CB-Dock2 following the standard procedure. Ligand-receptor docking results were evaluated visually for biochemical feasibility and docking results with the lowest Vina score were accepted. The predicted docking poses were evaluated using pymol3.7, and the predicted active site resides were identified for mutation.

### mPTER mutagenesis

A Q5 Site-Directed Mutagenesis Kit (NEB, E0554S) was used to introduce mutations in amino acid residues predicted to play a role in stabilizing zinc ions, interacting with N-acetyltaurine, or spatially constraining the active site of mPTER. The introduced mutations were subsequently verified through plasmid sequencing conducted by Genewiz.

### Activity-guided fractionation

6 kidneys from 10 to 14-week-old male C57BL/6J mice were homogenized homogenized using a Benchmark BeadBlaster Homogenizer at 4 °C. The cytosolic fraction was obtained using high-speed centrifugation at 15,000 rpm for 30 min at 4 °C. Then the mixture was concentrated using 3 kDa filter tubes (Millipore, UFC900324) by spinning down at 4,000 rpm for 1 h. The concentrated sample was diluted 50x into buffer containing 20 mM Tris pH 7.5 prior to anion exchange on a 1-ml HiTrap Q column (Cytiva, GE29-0513-25). The elution was performed from 0 mM to 500 mM NaCl in 20 mM Tris pH 7.5 over a gradient involving 30 column volumes. Following anion exchange, each fraction was evaluated for N-acetyltaurine synthesis and hydrolysis activities as described above. 3 fractions with the highest enzymatic activities were combined, concentrated and subjected to size exclusion on a Superose 6 Increase 10/300 GL column (Cytiva, GE29-0915-96). Each fraction from size exclusion was again evaluated for N-acetyltaurine synthesis and hydrolysis activity. The most active fraction was subjected to LC-MS analysis at the Vincent Coates Foundation Mass Spectrometry laboratory, Stanford University Mass Spectrometry.

### Shotgun proteomics

Samples were reduced with 10 mM dithiothreitol (DTT) for 20 minutes at 55 degrees Celsius, cooled to room temperature and then alkylated with 30 mM acrylamide for 30 minutes. They were then acidified to a pH ∼1 with 2.6 ul of 27% phosphoric acid, dissolved in 165 uL of S-trap loading buffer (90% methanol/10% 1M triethylammonium bicarbonate (TEAB)) and loaded onto S-trap microcolumns (Protifi, C02-micro-80). After loading, the samples were washed sequentially with 150 ul increments of 90% methanol/10% 100mM TEAB, 90% methanol/10% 20 mM TEAB, and 90% methanol/10% 5 mM TEAB solutions, respectively. Samples were digested at 47 °C for two hours with 600 ng of mass spectrometry grade Trypsin/LysC mix (Promega, V5113). The digested peptides were then eluted with two 35 µl increments of 0.2% formic acid in water and two more 40uL increments of 80% acetonitrile with 0.2% formic acid in water. The four elutions were consolidated in 1.5 ml S-trap recovery tubes and dried via SpeedVac (Thermo Scientific, San Jose CA). Finally, the dried peptides were reconstituted in 2% acetonitrile with 0.1% formic acid in water for LC-MS analysis.

Mass spectrometry experiments were performed using an Orbitrap Exploris 480 mass spectrometer (Thermo Scientific, San Jose, CA) attached to an Acquity M-Class UPLC system (Waters Corporation, Milford, MA). The UPLC system was set to a flow rate of 300 nl/min, where mobile phase A was 0.2% formic acid in water and mobile phase B was 0.2% formic acid in acetonitrile. The analytical column was prepared in-house with an I.D. of 100 microns pulled to a nanospray emitter using a P2000 laser puller (Sutter Instrument, Novato, CA). The column was packed with Dr. Maisch 1.9 micron C18 stationary phase to a length of approximately 25 cm. Peptides were directly injected onto the column with a gradient of 3-45% mobile phase B, followed by a high-B wash over a total of 80 minutes. The mass spectrometer was operated in a data-dependent mode using HCD fragmentation for MS/MS spectra generation.

RAW data were analyzed using Byonic v4.4.1 (Protein Metrics, Cupertino, CA) to identify peptides and infer proteins. A concatenated FASTA file containing Uniprot Mus musculus proteins and other likely contaminants and impurities was used to generate an *in silico* peptide library. Proteolysis with Trypsin/LysC was assumed to be semi-specific allowing for N-ragged cleavage with up to two missed cleavage sites. Both precursor and fragment mass accuracies were held within 12 ppm. Cysteine modified with propionamide was set as a fixed modification in the search. Variable modifications included oxidation on methionine, histidine and tryptophan, dioxidation on methionine and tryptophan, deamidation on glutamine and asparagine, and acetylation on protein N-terminus. Proteins were held to a false discovery rate of 1% using standard reverse-decoy technique. 247 proteins with at least 1 peptide match in total (**Supplemental Table 1**). PTER ranked #6 on the list.

### Preparation of mouse tissues for LC-MS analysis

50 µl plasma were mixed with 150 µl of a 2:1 mixture of acetonitrile:methanol and vortex for 30 s. The mixture was centrifuged at 15,000 rpm for 10 min at 4 °C and the supernatant was transferred to a LC–MS vial. For other mouse tissues, 50 µg samples were mixed with 150 µl of a 2:1 mixture of acetonitrile:methanol and homogenized using a Benchmark BeadBlaster Homogenizer at 4 °C. The mixture was spun down at 13,000 rpm for 10 min at 4 °C to pellet the insoluble materials. The supernatant was then transferred to a LC-MS vial.

### Measurements of metabolites by LC–MS

Metabolite measurements were performed using an Agilent 6520 Quadrupole time-of-flight LC– MS instrument as previously described^29^. MS analysis was performed using electrospray ionization (ESI) in negative mode. The dual ESI source parameters were configured as follows: the gas temperature was maintained at 250 °C with a drying gas flow of 12 l/min and the nebulizer pressure at 20 psi; the capillary voltage was set to 3,500 V; and the fragmentor voltage set to 100 V. The separation of polar metabolites was conducted using a Luna 5 μm NH2 100 Å LC column (Phenomenex 00B-4378-E0) with normal phase chromatography. Mobile phases were as follows: buffer A, 95:5 water:acetonitrile with 0.2% ammonium hydroxide and 10 mM ammonium acetate; buffer B, acetonitrile. The LC gradient initiated at 100% B with a flow rate of 0.2 ml/min from 0 to 2 min. The gradient was then linearly increased to 50% A/50% B at a flow rate of 0.7 ml/min from 2 to 20 min. From 20 to 25 min, the gradient was maintained at 50% A/50% B at a flow rate of 0.7 ml/min. N-acetyltaurine (Cayman, 35169) eluted around 12 min and taurine (sigma, T0625-500G) eluted around 13 min under the above conditions. The list of metabolites detected using LC-MS is summarized in **Supplemental Table 3**.

### General animal information

All animal experiments were performed according to protocols approved by the Stanford University Administrative Panel on Laboratory Animal Care. Mice were maintained in 12-h light–dark cycles at 22 °C and about 50% relative humidity and fed a standard irradiated rodent chow diet. Where indicated, a high-fat diet (D12492, Research Diets 60% kcal from fat) was used. Male C57BL/6J (stock number 000664), male C57BL/6J DIO mice (stock number 380050) and male MC4R-KO mice (stock number 032518) were purchased from the Jackson Laboratory. Whole-body PTER-KO mice (catalogue number C57BL/6N(Jax)-Pter^em1^(IMPC)^Bay^) were obtained from the Baylor KOMP2 group of International Mouse Phenotyping Consortium (IMPC). For intraperitoneal injections of mice with compounds, compounds were dissolved in saline (Teknova, S5825). Compounds were administered to mice every day by intraperitoneal injections at 10 μl/g body weight at the indicated doses. For chronic intraperitoneal injection, oral gavage and subcutaneous injection experiments, mice were mock treated with saline for 3 to 5 days until body weights were stabilized. For control IgG or anti-GFRAL antibody treatment, mice were subcutaneously injected with 10 mg/kg antibodies once every 3 days. For GLP-1 and Exendin-3 injection, GLP-1 and Exendin-3 powder was first dissolved in 18:1:1 saline:DMSO:kolliphore and then injected (GLP-1: 2 mg/kg/day, IP; Exendin-3, 0.1 mg/kg/day, IP). Unless specified, compounds were administered around 6 pm. For measuring known feeding-regulating polypeptide hormones, blood plasma was collected at 9 am and ELISA kits were used following manufacturer’s instructions (Leptin: Crystal Chem, 90030; GLP-1: Sigma, EZGLP1T-36K; GDF-15: R&D Systems, MGD150; Adiponectin: Crystal Chem, 80569; FABP4: Novus biologicals, NBP2-82410; Insulin: Crystal Chem, 90080; ALT: Cayman, 700260; AST: Cayman, 701640; triglycerides: Cayman, 10010303).

### Breeding and genotyping of PTER-KO mice

PTER-KO and WT animals were generated through heterozygous breeding crosses and weaned around postnatal day 21. Genotyping was performed using the following procedures: tail clippings were collected from littermates and boiled for 30 min at 95 °C in 100 μl of 50 mM NaOH to extract genomic DNA. The solution was neutralized by adding 42 μl of 0.5 M Tris (pH 7.5). PCRs were performed by using primers for either the PTER WT allele (forward, 5′-TCATGTCCCACCTTGACAGGTAAGCGGGTC-3′; reverse, 5′-CAGTTGTAGCAGCCATGAACACTATTGTGC-3′) or PTER KO allele (forward, 5′-GGGTAATATACTTGTCAAACCATGCT-3′; reverse, 5′-CAGTTGTAGCAGCCATGAACA-3′). Promega GoTaq master mix (Promega, PRM7123) was used for the PCR reaction. Each 25 μl reaction consisted of 12.5 μl of the Promega master mix, 2.5 μl of a 10 μM mixture of forward and reverse primers, 2 μl of genomic DNA and 8 μl of ultrapure water. The thermocycling program on a Bio-Rad C1000 Touch Thermo Cycler began with an initial 90 s at 98 °C, followed by cycles of 30 s at 98 °C, 30 s at 58 °C for KO primers and 50 °C for WT primers and 30 s at 72 °C, followed by 5 min at 72 °C and finally held at 4 °C. PCRs for WT primers consisted of 41 cycles, whereas PCRs for KO primers consisted of 35 cycles. Samples were run on a 1.5% agarose gel with 0.1 mg/ml ethidium bromide. WT alleles are expected to yield a PCR product of 699 base pairs in size whereas KO alleles are expected to yield PCR products that are 479 base pairs in size.

### Taurine water supplement

2.5% (w/v) taurine (sigma, T0625-500G) was dissolved in mouse drinking water and supplemented to 4-week-old male PTER-KO and WT mice. Taurine water was freshly prepared every 3 days while mice were on a high-fat diet (D12492, Research Diets 60% kcal from fat). Body weights, food intake and water consumption were measured every 3 days. No adverse effects were observed in mice fed with taurine water.

### N-aectyltaurine ex vivo kinetic analysis

Kidneys from 8-week-old PTER-KO and WT mice were dissected out and incubated with 9x (v/w) pre-warmed Williams Medium E (Quality Biological, 112-033-101) supplemented with 5 µM heavy N-acetyltaurine (Acme) at 37 °C on a shaker. 30 µl supernatant media was collected at 0, 15, 30, 45, 60, 90, 120 and 240 min of incubation. Metabolites were extracted and analyzed by LC-MS as previously described.

### Adipose lipolysis in vivo and ex vivo

Blood plasma and epidydimal fat were collected from 4-month-old male DIO C57BL/6J mice receiving saline, N-acetyltaurine (NAT, 15 mg/kg, IP) or norepinephrine (NE, 0.5mg/kg, IP) treatment. Blood glycerol contents were determined using a glycerol quantification kit (Sigma, F6428-40ML). For mature adipocyte lipolysis ex vivo, epidydimal fat from 4-month-old male DIO C57BL/6J mice was dissected out and dissociated using 2 mg/ml Collagenase B (Worthington, CLSAFB) and 1 mg/ml soybean trypsin inhibitor (Worthington, LS003570). Digested adipose tissues were spun down at 500 g for 3 min to isolate the floating layer of mature adipocytes. 1 million mature adipocytes were collected and incubated with saline, 50 μM N-acetyltaurine (NAT) or 1 μM norepinephrine (NE) at 37 °C on a shaker for 1 h. Then released glycerol was determined using a glycerol quantification kit (Sigma, F6428-40ML).

### Indirect calorimetry and physiological measurements

8-to 9-week-old male PTER-KO and WT mice (N=9/group) were supplemented with 2.5% (w/v) taurine water and fed on a high-fat diet for 4 weeks. Taurine water was freshly prepared every 3 days when body weights and food intake were measured. Before the body weights of PTER-KO mice started to be significantly different from WT mice (4 weeks on taurine water), metabolic parameters including oxygen consumption, carbon dioxide production, respiratory exchange ratio (RER), food intake and ambulatory movement of mice were measured using the environment-controlled home-cage CLAMS system (Columbus Instruments) at the Stanford Diabetes Center. A separate cohort of 12-to 13-week-old male PTER-KO and WT mice (N=8/group) were supplemented with 2.5% (w/v) taurine water and fed on a high-fat diet for 8 weeks before putting into the metabolic cages for analysis. Mice were housed in the metabolic chambers for 36 h prior to the start of the experiment. Data collected during a complete 24-hour day-night cycle were used for analysis. Energy expenditure calculations were normalized for body weight. P-values were calculated from two-tailed unpaired t-tests.

### Mouse exercise training protocols

A Columbus Instrument animal treadmill with six lanes (Columbus, 1055-SRM-D65) was employed for the treadmill running experiments. Prior to commencing the treadmill running, mice were given a 5-minute acclimation period. The initial treadmill running phase began at a speed of 7.5 m/min with a 4° incline, following the procedure as previously described^29^. At intervals of 3 minutes, both the speed and incline were incrementally increased by 2.5 m/min and 2°, respectively. Once the maximum parameters of 40 m/min in speed and a 30° incline were attained, they remained constant until the mice reached a state of exhaustion, defined as when the mice remained on the shocker at the rear of the treadmill for longer than 5 seconds. PTER-KO and WT mice were exercised every other day, while on a high-fat diet (60% kcal from fat) for a whole duration of 6 weeks. Running was performed in the mid-morning for all experiments. Body weights and food intake were measured right before each exercise training session.

### Glucose tolerance and insulin tolerance tests in mice

For glucose tolerance tests, mice were fasted for 6 h (fasting starting 7 a.m. in the morning) and then intraperitoneally injected with glucose at 2 g/kg body weight. Blood glucose levels were measured at 0, 20, 40, 60, and 120 min via tail bleeding using a glucose meter. For insulin tolerance tests, mice were fasted for 6 h (fasting starting 7 a.m. in the morning) and then intraperitoneally injected with insulin in saline 0.75 U/kg body weight. Blood glucose levels were measured at 0, 20, 40, 60, and 120 min via tail bleeding using a glucose meter.

### hCom2 bacterial strains and culture conditions

Individually cultivated hCom2 strains were obtained from the Microbiome Therapies Initiative (MITI). All strains were cultured in one of two growth media: mega medium (MM) and chopped meat medium w/ rumen fluid and carbohydrates (CMM). Cultures were incubated at 37 °C in an anaerobic chamber (Coy Laboratories) in an atmosphere of 5% hydrogen, 10% CO2 and 85% N2. Cultures were stored in anaerobically prepared 25% glycerol/water (v/v). All medium and reagents used in the anaerobic chamber were pre-reduced for at least 48 h.

### Synthetic community construction

Frozen stocks in 96-well plate matrix tubes were thawed, and 300 µl of each thawed culture was used to inoculate 40 ml of growth medium in 50 ml falcon tubes. After 72 hr, non-normalized cultures of all strains were pooled into a mixture. A 1 ml aliquot of the resulting mixed culture was stored at −80 °C for metagenomic sequencing, The remainder of the mixed culture was subjected to centrifugation (4700 x g, 30 min). The cell pellet was washed with an equal volume of pre-reduced sterile phosphate-buffered saline (PBS), and then resuspended in 1/120 of the initial volume of 25% glycerol/water (v/v) solution. Aliquots of the resulting synthetic community were stored in 2 ml cryovials (Corning, 430659) at −80 °C until use.

### Gnotobiotic mouse experiments

Germ free C57BL/6N mice (male, 6-8 weeks of age) were originally obtained from Taconic Biosciences (Hudson, NY) and colonies were maintained in gnotobiotic isolators and fed ad libitum. The Institutional Animal Care and Use Committee(IACUC) at Stanford University approved all procedures involving animals. Glycerol stocks of synthetic communities were thawed and shaken well at room temperature, and mice were orally gavaged with 200 µl of the mixed culture. To ensure efficient colonization by all strains in the community, mice were gavaged using the same procedure twice on different days for all experiments. Mice were fed standard chow (LabDiet, 5k67), fresh fecal pellets were collected weekly at the same time of day and stored at −80 °C prior to analysis. The mice were maintained on a standard diet (LabDiet, 5k67; 0.2% Trp) for 4 weeks before sacrifice (fed ad libitum). Fresh fecal samples from GF mice and hCom2-colonized mice were collected, normalized by weight, homogenized, and spun down to isolated live bacteria for in vitro incubation. Mice were euthanized humanely by CO_2_ asphyxiation and the plasma were collected in a BD blood tube (BD 365967) and stored on ice. Plasma samples were centrifuged at 16,000 x g for 20 min, and supernatant were stored in −80 °C until use.

### hCom2 in vitro screening

Individually cultivated hCom2 strains were resuspended in a standard amino acid complete (SAAC) medium as previously described^37^. 100 µl of cell suspension from each strain was incubated with 10 mM taurine and 10 mM acetate in 300 µl SAAC medium in an anaerobic chamber (Coy Laboratories) in an atmosphere of 5% hydrogen, 10% CO2 and 85% N2. Cells were spun down after 48hr incubation to obtain cell pellets and conditioned medium. Metabolites were extracted and analyzed by LC-MS. OD600 prior to and after incubation was measured.

### Antibiotic treatment in mice

12 to 14-week-old mice were treated with antibiotic mixture mixture (chloramphenicol (Sigma, C0378-25G), spectinomycin dihydrochloride pentahydrate (Sigma, S4014-25G), apramycin sulfate (Sigma, A2024-5G), tetracycline hydrochloride (Sigma, T7660-5G), kanamycin (ThermoScientific, 11815032) and ampicillin (Sigma, A9518) at 1g/L per antibiotic) was administered in drinking water ad libitum and orally gavaged (0.5 ml) every other day for a whole duration of 2 weeks. Before blood was collected from these mice, fresh fecal samples were collected using sterile pre-weighted Eppendorf tubes and labeled with unique identifiers. Samples were immediately stored at −80°C until further processing. Fecal samples were normalized by weight, homogenized, and filtered prior to DNA extraction. DNA was extracted from fecal samples using the Qiagen Mini Prep Kit following the manufacturer’s protocol. Extracted DNA was stored at −20 °C until qPCR analysis.

Universal bacterial primers targeting the V3 region of the bacterial 16S rRNA were selected (forward HV3-16S primer 5′CCAGACTCCTACGGGAGGCAG-3′ and the reverse HV3-16S primer 5′-CGTATTACCGCGGCTGCTG-3′). Mouse genomic DNA was used as house-keeping gene for qPCR analysis. All reactions were carried out with 10ng total DNA and SsoAdvanced Universal SYBR Green Supermix (BioRad, 1725274) in CFX Opus 384. Reactions were held at 95 °C for 10 minutes, followed by 40 cycles of 95 °C for 15 s and 60 °C for 60 s. The number of 16s DNA copies was subsequently determined and normalized to the number of mouse genomic DNA copies in the same fecal sample.

## Data availability

All data generated or analyzed during this study are included in this published article and its supplementary information files. Source data are provided with this paper.

## Code availability

No new code is generated in this study.

## Acknowledgments

We thank members of the Long lab for helpful discussions. We gratefully acknowledge the staff at the Baylor KOMP2 group of International Mouse Phenotyping Consortium (IMPC) for the production and shipment of the Cas9-RGN null allele Pter^em1^(IMPC)^Bay^ transgenic mice, Dr. Paul Emmerson (Eli Lilly & Co.) for sharing the anti-GFRAL neutralizing antibody and control IgG antibody, and Microbiome Therapies Initiative (MITI) for providing individually cultivated hCom2 strains. This work was supported by the US National Institutes of Health (DK105203 and DK124265 to J.Z.L., DK111916 to K.J.S), the Stanford Diabetes Research Center (P30DK116074), the Stanford Cardiovascular institute (CVI), the Weintz Family COVID-19 research fund (K.J.S), American Heart Association (AHA), the Stanford School of Medicine, the Jacob Churg Foundation (K.J.S), the McCormick and Gabilan Award (K.J.S), the Wu Tsai Human Performance Alliance (research grant to J.Z.L, postdoctoral fellowship to X.L. and M.D.M.G.), the Stanford Diabetes Research Center (P30DK116074 to J.Z.L.), the Phil & Penny Knight Initiative for Brain Resilience at the Wu Tsai Neurosciences Institute (research grant to J.Z.L.), the American Heart Association (postdoctoral fellowship 24POST1196199 to W.W, 905674 to M.Z.), K99/R00 NIH Pathway to Independence Award (K99AR081618 to M.Z.), and Stanford Maternal Research Institute (postdoctoral fellowship to D.X.).

## Contributions

Conceptualization, W.W. and J.Z.L.; methodology, W.W.; investigation, W.W., X.L., A.L.M, S.F., R.E.M., P.E.C, X.Z., J.R., N.L., S.X., M.Z., M.D.M.G., S.D.T, J.C.C., L.W.W., S.C.R, L.C., D.X., F.S., W.H., C.B.R., and C.J.; writing – original draft, W.W. and J.Z.L.; writing – review & editing, W.W. and J.Z.L.; resources, K.J.S., C.J., M.A.F., J.Z.L.; supervision and funding acquisition, J.Z.L.

## Ethics declarations

### Competing interests

A provisional patent application has been filed by Stanford University on PTER/N-acetyltaurine for the treatment of cardiometabolic disease.

## Extended Figure Legends

**Extended Data Fig. 1.**
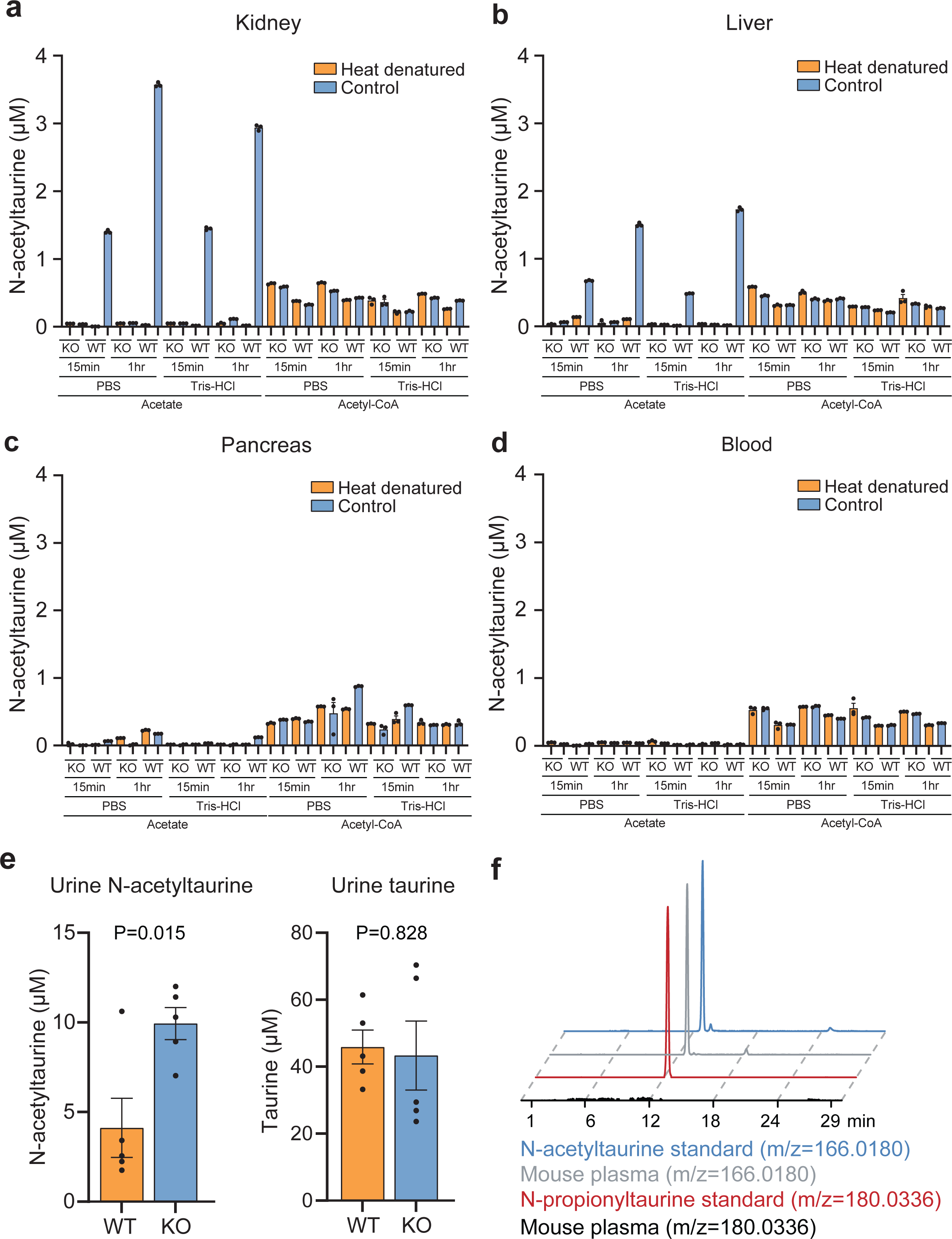
Additional biochemical characterization of PTER-KO mice. Related to Fig. 3. (a-d) Production of N-acetyltaurine from cell lysates of kidney (a), liver (b), pancreas (c) or blood plasma (100 µg) following incubation at 37°C under the indicated buffer condition and incubation time. The following concentrations of substrates were used: taurine (10 mM), acetate (10 mM), acetyl-CoA (100 µM). N=3/condition. (e) Urine N-acetyltaurine concentration (left) and taurine concentration (right) of 13- to 14-week-old male WT or PTER-KO mice. N=5/group. (f) Representative extracted ion chromatograms of synthetic N-acetyltaurine and N-propionyltauirne standards and endogenous peaks of blood plasma from 14-week old wild-type male mice. Data are shown as mean ± SEM. In (e), P-values were calculated from two-tailed unpaired t-tests.

**Extended Data Fig. 2.**
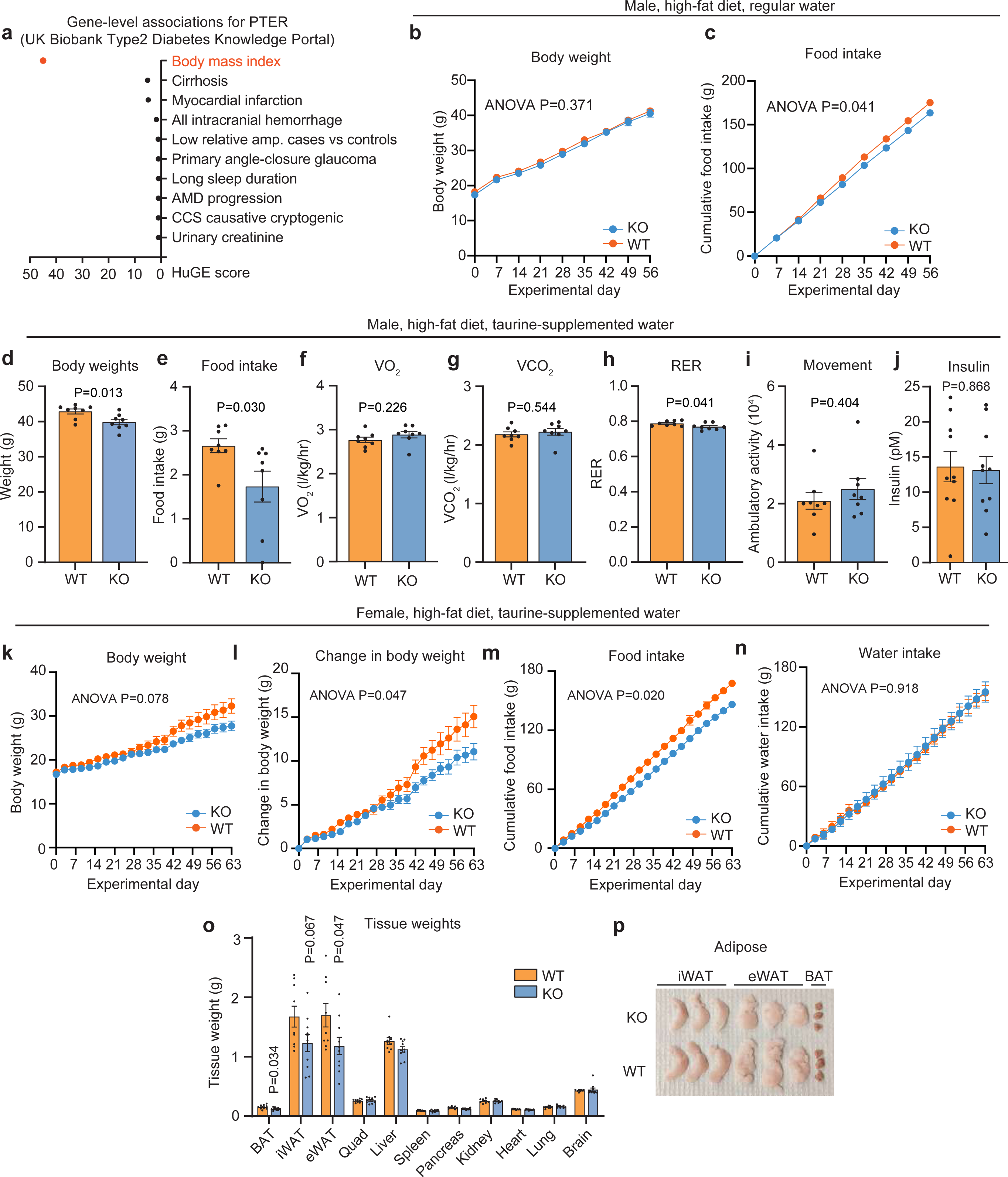
Additional metabolic characterization of male and female PTER-KO mice on high fat diet and taurine-supplemented water. Related to Fig. 4. (a) Human genetic evidence (HuGE) score of phenotype associations for the *PTER* gene locus from the Type 2 Diabetes Knowledge Portal. (b,c) Body weight (b) and food intake (c) of 12- to 13-week-old male PTER-KO mice (N=10, blue line) or WT mice (N=14, orange line) subjected to a high-fat diet feeding alone for a period of 8 weeks. (d-j) Metabolic chamber analysis of 12- to 13-week-old-male WT or PTER-KO mice after 8 weeks of high fat diet/taurine water supplementation (2.5% w/v). N=8/group. RER, respiratory exchange ratio. (k-p) Body weight (k), change in body weight (l), cumulative food intake (m), and water intake (n), tissue weights (o) and representative adipose tissues (p) of 13- to 14-week-old female WT or PTER-KO mice on high fat diet and after taurine water supplementation (2.5% w/v). N=10/group. iWAT, inguinal white adipose tissue; eWAT, epididymal white adipose tissue; BAT, brown adipose tissue; Quad, quadricep muscles. Data are shown as mean ± SEM. In (b,c) and (k-n), P-values were calculated from two-way ANOVA with post hoc Sidak’s multiple comparisons test. In (d-j) and (o), P-values were calculated from two-tailed unpaired t-tests.

**Extended Data Fig. 3.**
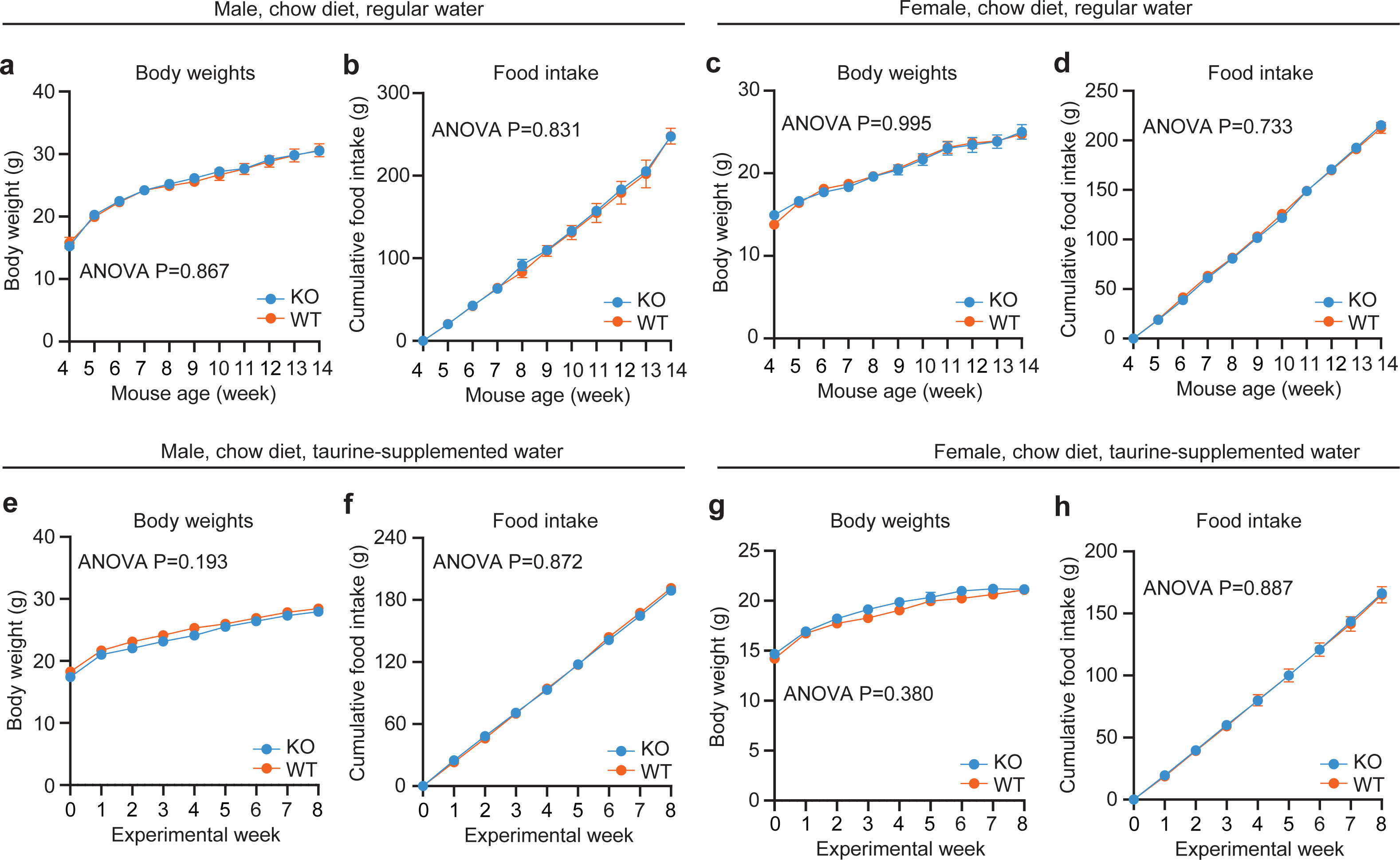
Additional metabolic characterization of chow-fed male and female PTER-KO mice. Related to Fig. 4. (a-h) Body weight (a,c,e,g) and cumulative food intake (b,d,f,h) of chow-fed 12- to 14-week-old male mice (a,b,e,f) or female mice (c,d,g,h) with regular water (a-d) or with 2.5% w/v taurine-supplemented water (e-h). Data are shown as mean ± SEM. P-values were calculated from two-way ANOVA with post hoc Sidak’s multiple comparisons test.

**Extended Data Fig. 4.**
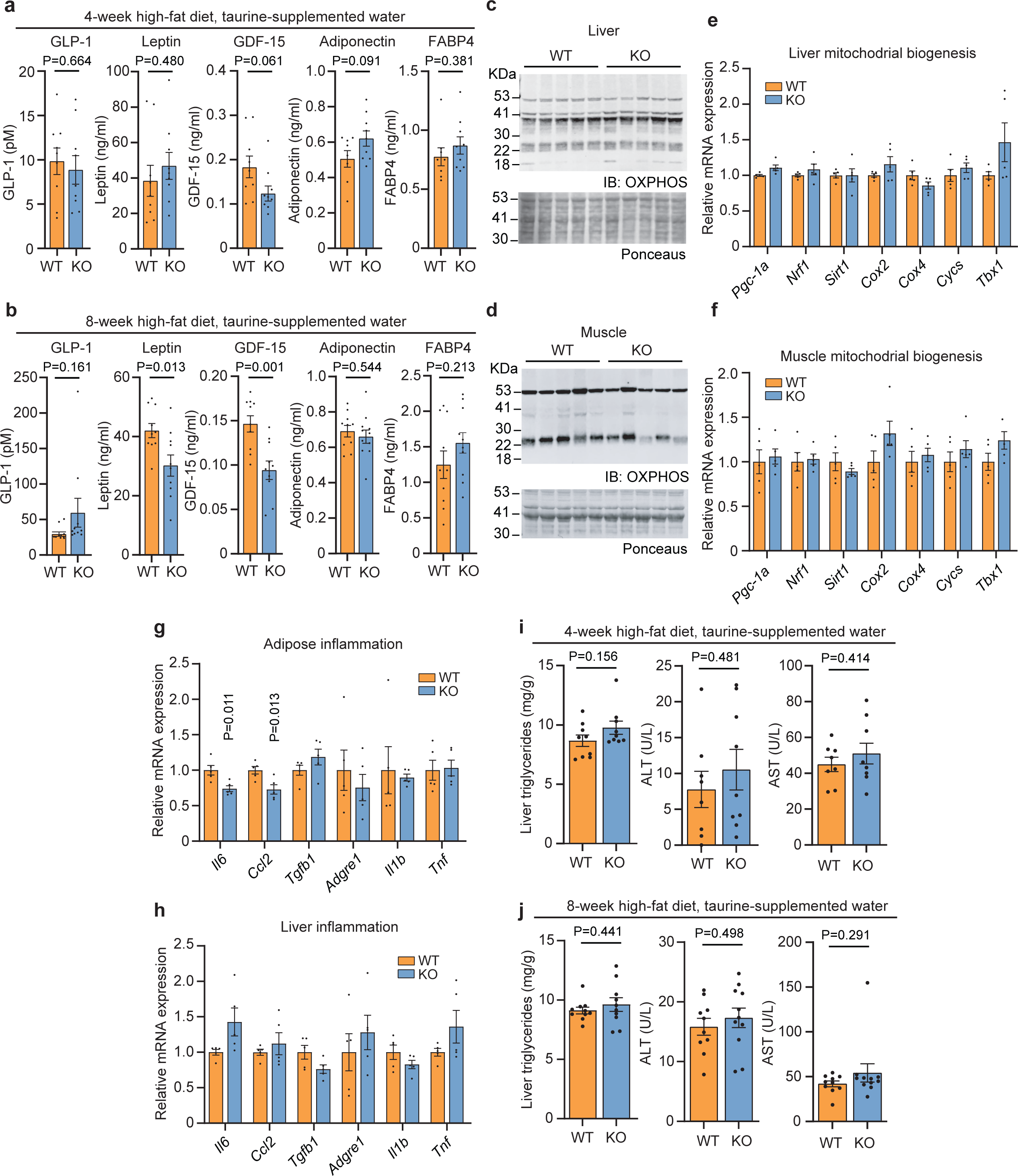
Additional molecular characterization of PTER-KO mice. Related to Fig. 4. (a,b) Blood plasma levels of the indicated hormones from 8 to 9-week-old-male WT or PTER-KO mice after 4 weeks of high fat diet/taurine water supplementation (2.5% w/v) (a) or from 12- to 13- week-old-male WT or PTER-KO mice after 8 weeks of high fat diet/taurine water supplementation (2.5% w/v) (b). In (a), N=8/group. In (b), N=10/group. (c,d) Western blotting with anti-OXPHOS cocktail antibody (top) and Ponceaus stain (bottom) of liver (c) and quadricep muscles (d) from 12- to 13-week-old-male WT or PTER-KO mice after 8 weeks of high fat diet/taurine water supplementation (2.5% w/v). (e-h) mRNA expression of indicated genes in liver (e,h), quadricep muscles (f), or epididymal white adipose tissue (g) from 12 to 13-week-old-male WT or PTER-KO mice after 8 weeks of high fat diet/taurine water supplementation (2.5% w/v). N=5/group. (i,j) Liver triglycerides and liver enzymes from 8- to 9-week-old-male WT or PTER-KO mice after 4 weeks of high fat diet/taurine water supplementation (2.5% w/v) (l) or 12- to 13-week-old-male WT or PTER-KO mice after 8 weeks of high fat diet/taurine water supplementation (2.5% w/v) (j). In (i), N=8/group. In (j), N=10/group. Data are shown as mean ± SEM. P-values were calculated from two-tailed unpaired t-tests.

**Extended Data Fig. 5.**
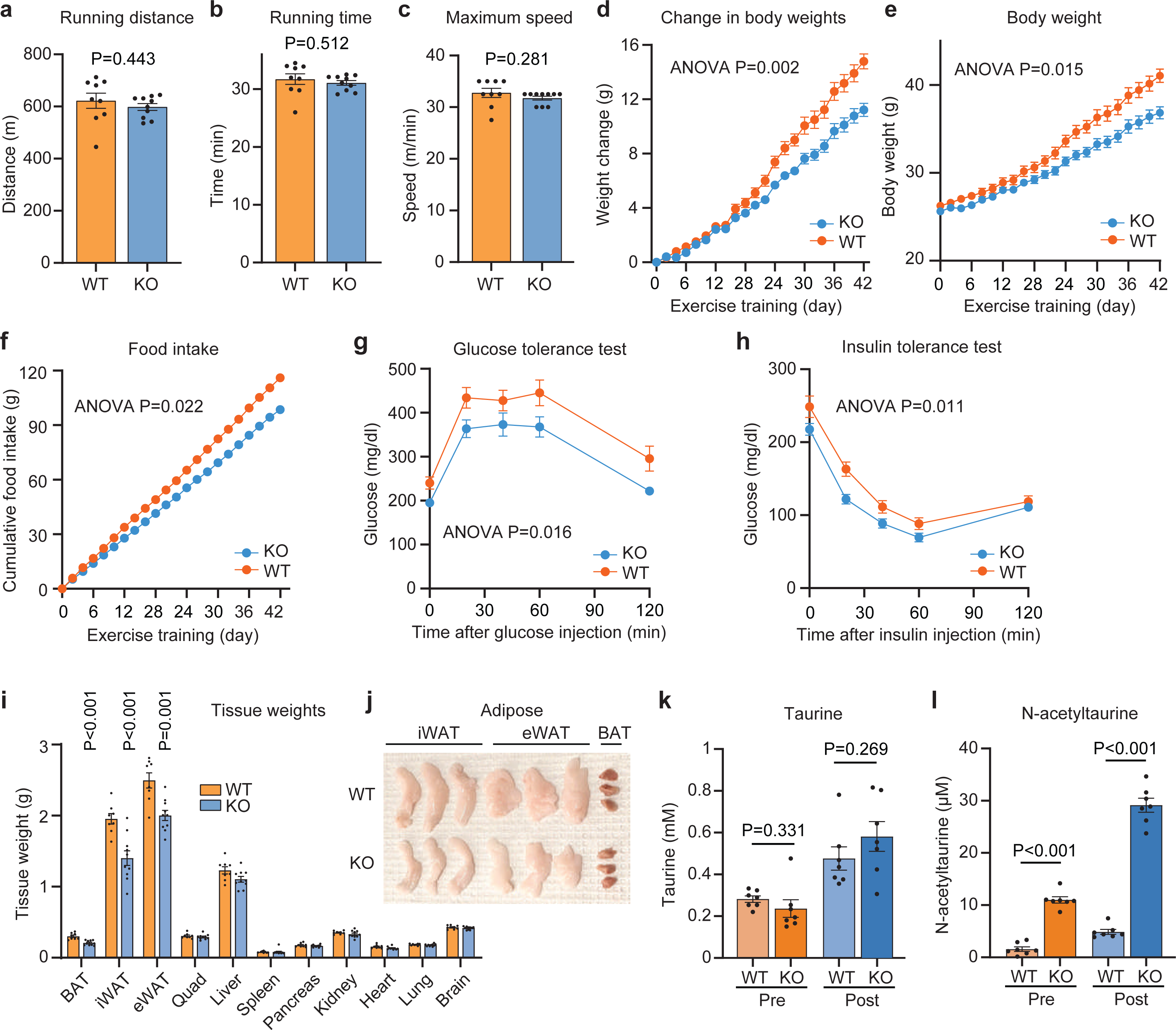
Metabolic phenotype of male PTER-KO mice on a chronic treadmill exercise training protocol. Related to Fig. 4. (a-c) Running distance (a), running time (b) and maximum speed (c) of 10- to 11-week-old male PTER-KO mice (N=10, blue line and box) or WT mice (N=9, orange line and box) following a single bout of treadmill exercise running until exhaustion. (d-l) Change in body weight (d), body weight (e), cumulative food intake (f), glucose tolerance test (g), insulin tolerance test (h), tissues weights (i), representative adipose tissues (j), plasma taurine levels (k), and plasma N-acetyltaurine levels (l) of 10- to 11-week-old male PTER-KO mice (N=10, blue line and box) or WT mice (N=8, orange line and box) after a 6-week treadmill exercise training protocol (see **Methods**). Data are shown as mean ± SEM. In (a-c) and (i-l), P-values were calculated from two-tailed unpaired t-tests. In (d-h), P-values were calculated from two-way ANOVA with post hoc Sidak’s multiple comparisons test.

**Extended Data Fig. 6.**
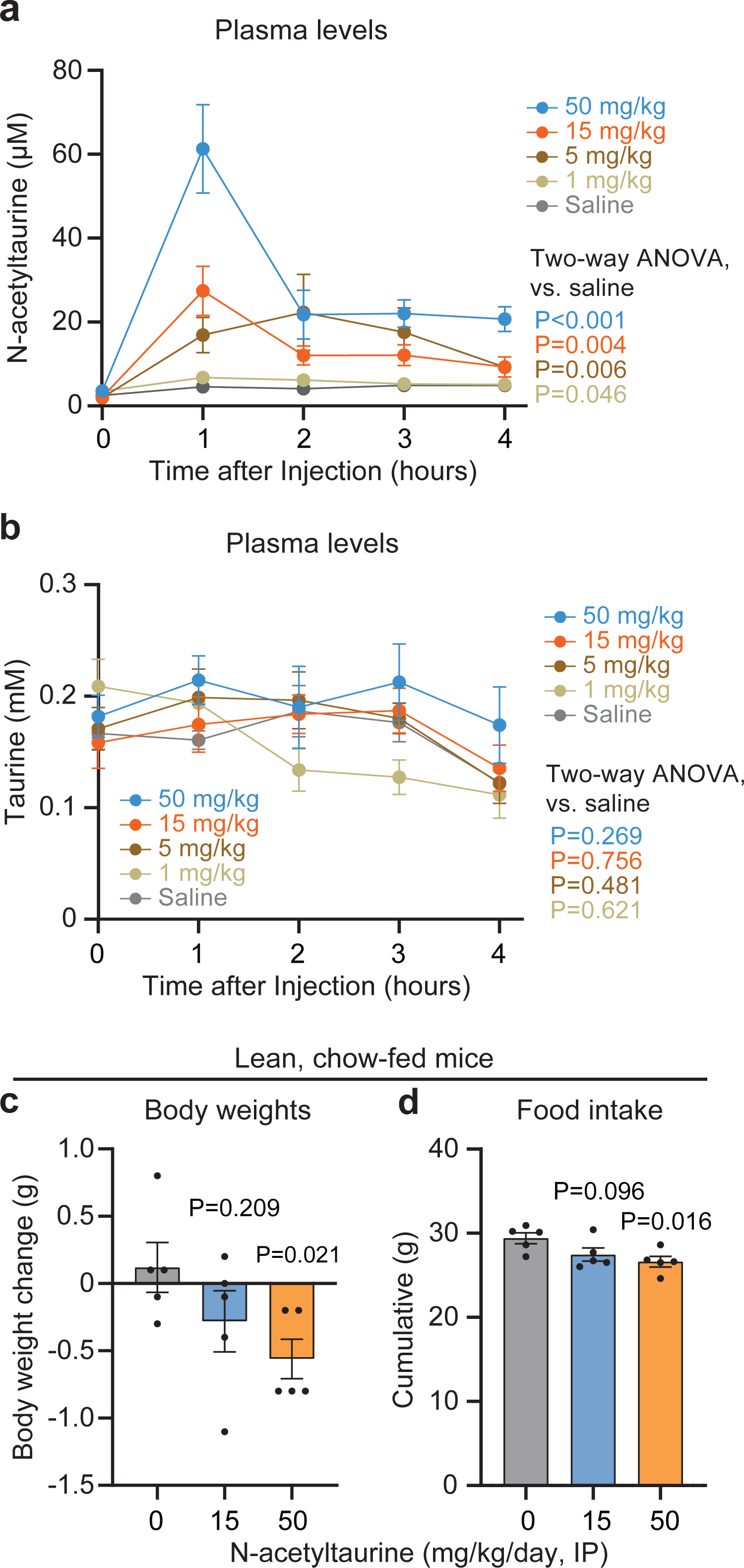
Additional characterization of N-acetyltaurine administration to wild-type mice. Related to Fig. 5. (a,b) Blood plasma concentrations of N-acetyltaurine (a) or taurine (b) from 26-to 28-week-old male DIO C57BL/6J mice after intraperitoneal injection with indicated doses of N-acetyltaurine. N=5/group. (c-d) Body weight change (c) and cumulative food intake (d) of 13-week-old chow-fed male C57BL/6J mice following 7 days of treatment with the indicated dose of N-acetyltaurine (intraperitoneal injection). N=5/group. Data are shown as mean ± SEM. In (c,d), P-values were calculated from two-tailed unpaired t-tests. In (a,b), P-values were calculated from two-way ANOVA with post hoc Sidak’s multiple comparisons test.

**Extended Data Fig. 7.**
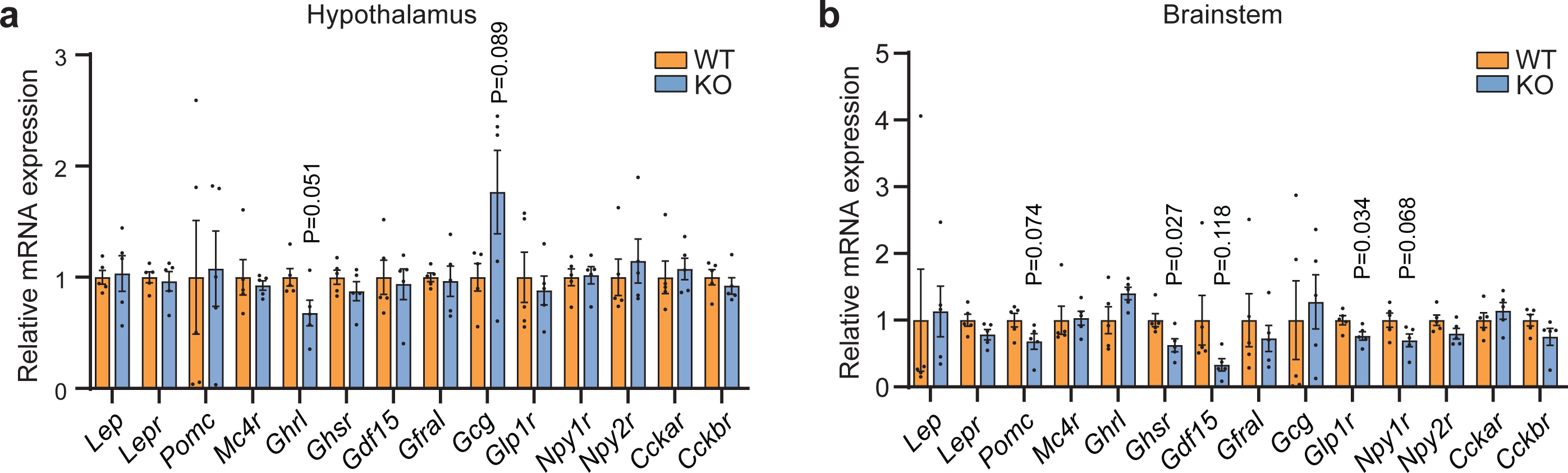
Gene expression in hypothalamus and brainstem from PTER-KO mice. Related to Fig. 5. (a,b) mRNA expression of indicated genes of hypothalamus (a) and brainstem (b) from 12-to 13-week-old-male WT or PTER-KO mice after 8 weeks of high fat diet/taurine water supplementation (2.5% w/v). N=5/group. Data are shown as mean ± SEM. P-values were calculated from two-tailed unpaired t-tests.

**Extended Data Fig. 8.**
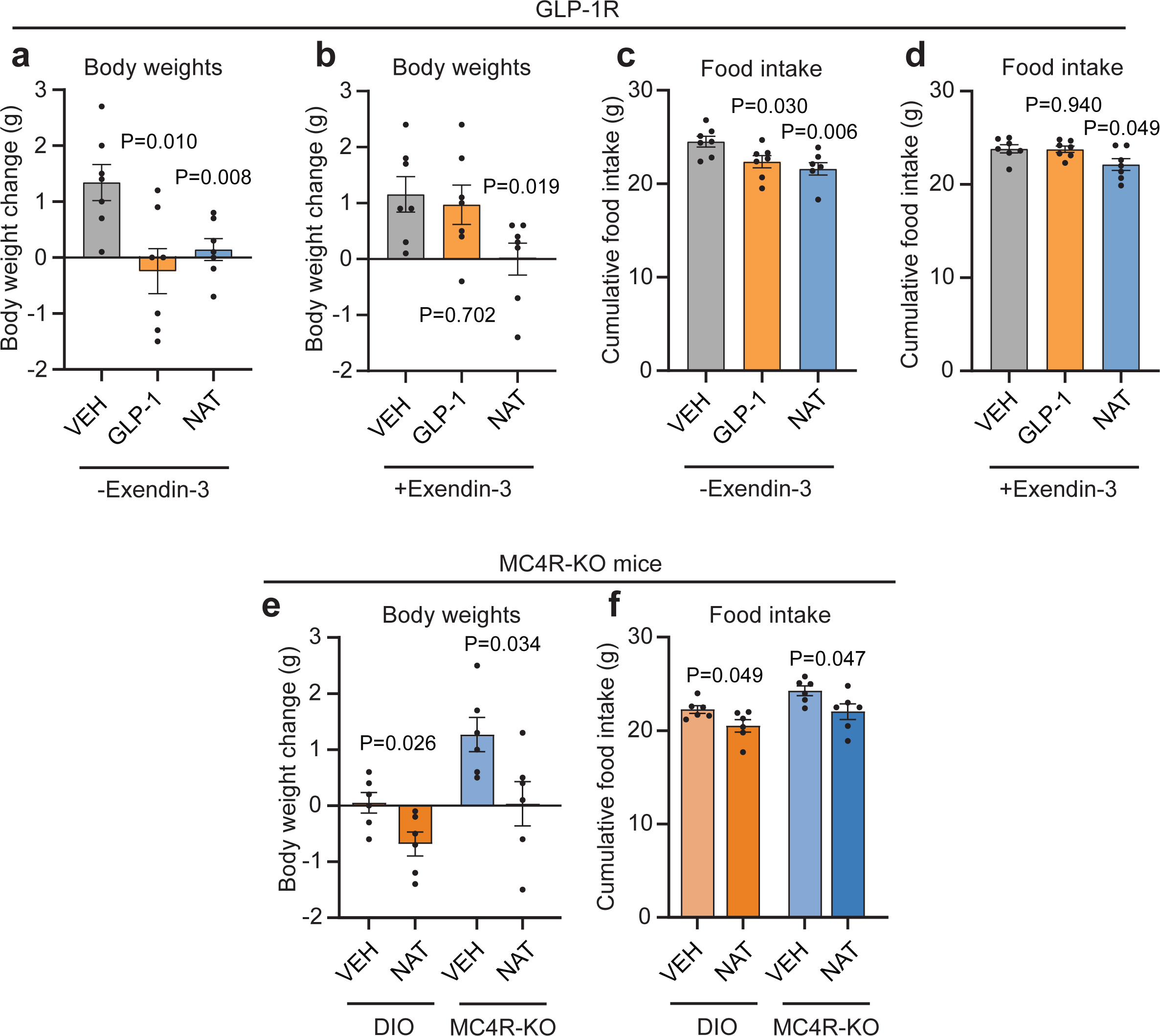
Requirement for GLP1R and MC4R in the anorexigenic effects of N-acetyltaurine. Related to Fig. 5. (a-d) Body weight (a,b) and cumulative food intake (c,d) of 6 to 7-month-old DIO male C57BL/6J mice after a 7-day treatment of saline or N-acetyltaurine (15 mg/kg/day, IP) or GLP-1 (2 mg/kg/day, IP) with or without Exendin-3 (0.1 mg/kg/day, IP). N=7/group. NAT, N-acetyltaurine. (e,f) Body weight (e) and cumulative food intake (f) of 5-month-old DIO male C57BL/6J mice or 3 to 4-month-old MC4R-KO mice on high fat diet after a 7-day treatment of saline or N-acetyltaurine (15 mg/kg/day, IP). N=6/group. NAT, N-acetyltaurine. Data are shown as mean ± SEM. P-values were calculated from two-tailed unpaired t-tests.

**Extended Data Fig. 9.**
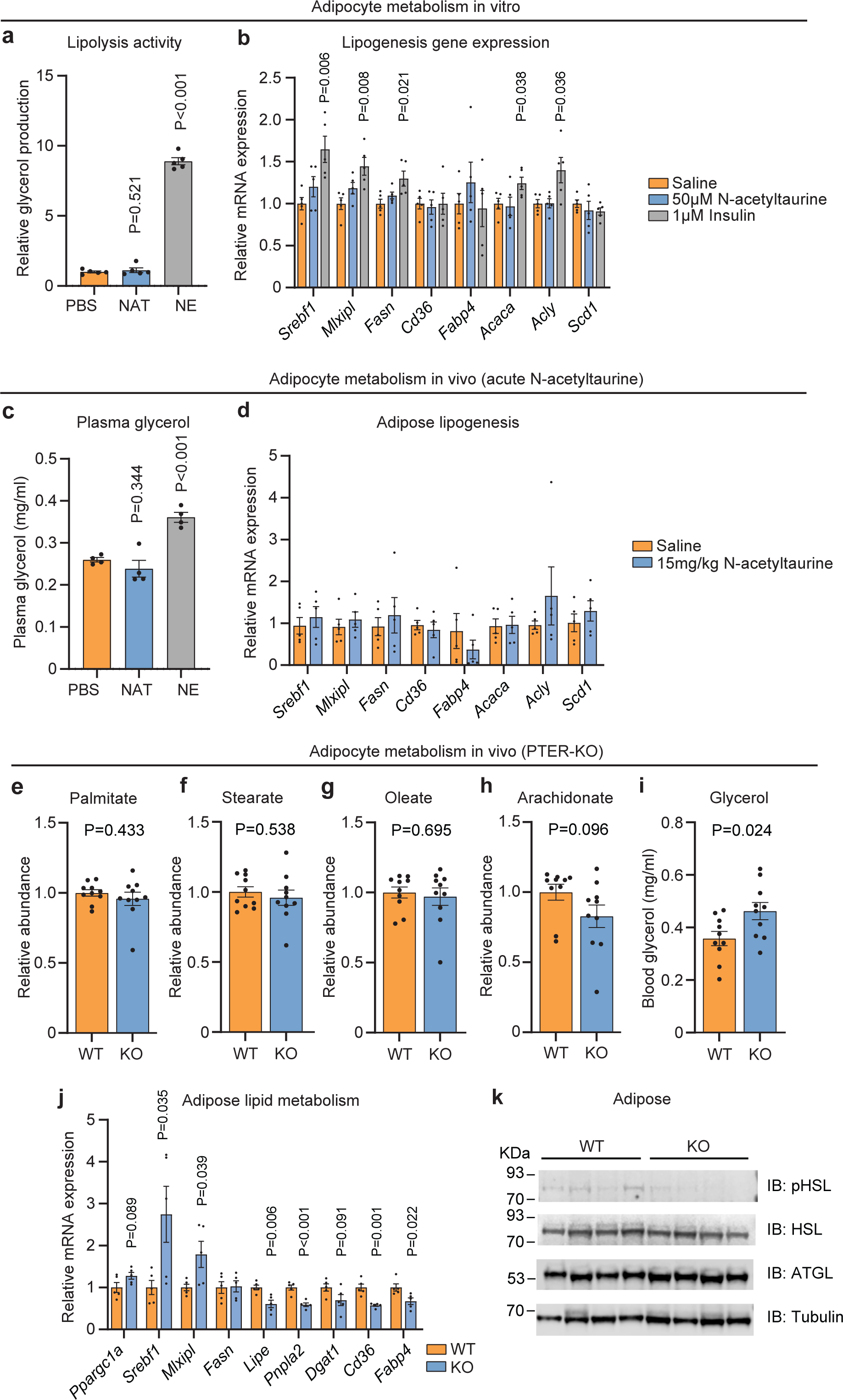
Direct and indirect effects of N-acetyltaurine on adipose. Related to Fig. 5. (a) Glycerol production from mature epididymal adipocytes isolated from 4-month-old male C57BL/6J mice after incubation with 50 μM N-acetyltaurine (NAT) or 1 μM norepinephrine (NE) at 37 °C for 1 h. N=5/group. (b) mRNA expression of indicated genes from mature epididymal adipocytes isolated from 4-month-old male C57BL/6J mice and incubated with PBS or 50 μM N-acetyltaurine or 1 μM insulin at 37 °C for 4 h with constant shaking. N=5/group. (c) Plasma glycerol levels of 4-month-old male DIO C57BL/6J mice one hour after a single administration of N-acetyltaurine (NAT, 15 mg/kg, IP) or norepinephrine (NE, 0.5mg/kg, IP) treatment. N=4/group. (d) mRNA expression of indicated genes from epididymal white adipose tissues of 4-month-old male DIO C57BL/6J mice four hours after a single administration of N-acetyltaurine (15 mg/kg, IP) treatment. N=5/group. (e-k) Plasma palmitate (e), stearate (f), oleate (g), arachidonate (h) and glycerol (i), mRNA expression of indicated genes from epididymal white adipose tissue (j), Western blotting of epididymal white adipose tissues with the indicated antibodies (k) of 13 to 14-week-old male WT or PTER-KO mice after 8 weeks on high fat diet and taurine water supplementation (2.5% w/v). In (e-i), N=10/group. In (j), N=5/group. In (k), N=4/group. Data are shown as mean ± SEM. In (a-j), P-values were calculated from two-tailed unpaired t-tests.

**Extended Data Fig. 10.**
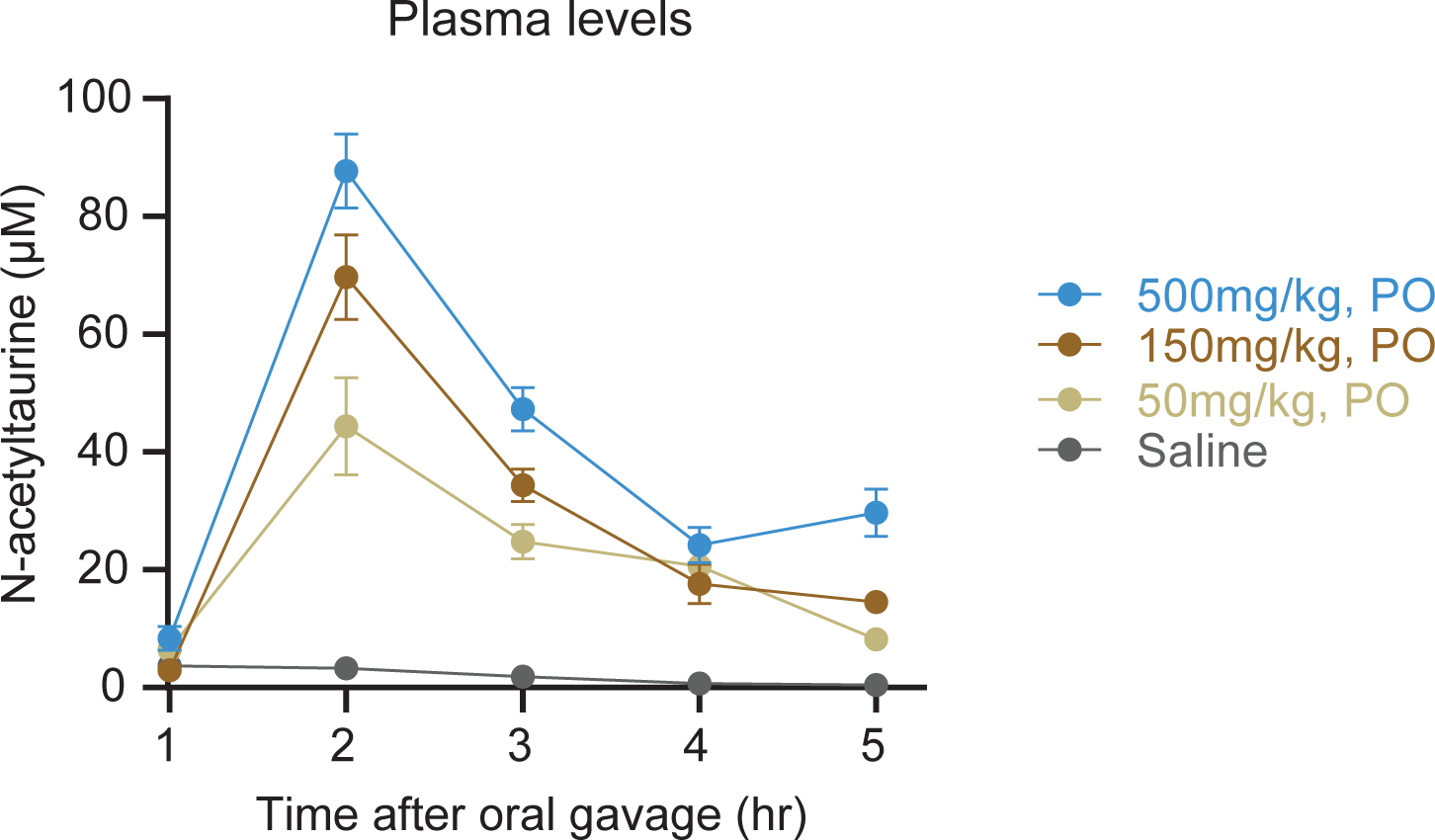
Pharmacokinetics of oral N-acetyltaurine administration to DIO mice. Related to Fig. 6. Blood plasma concentrations over time of N-acetyltaurine from 7-month-old male DIO C57BL/6J mice after oral gavage with indicated doses of N-acetyltaurine. N=5/group.

## References

1 Lourenço, R. & Camilo, M. E. Taurine: a conditionally essential amino acid in humans? An overview in health and disease. Nutr Hosp 17, 262–270 (2002).

2 Ripps, H. & Shen, W. Review: taurine: a “very essential” amino acid. Mol Vis 18, 2673–2686 (2012).

3 Lambert, I. H., Kristensen, D. M., Holm, J. B. & Mortensen, O. H. Physiological role of taurine--from organism to organelle. Acta Physiol (Oxf) 213, 191–212 (2015). 10.1111/apha.12365

4 Stipanuk, M. H. Metabolism of sulfur-containing amino acids. Annu Rev Nutr 6, 179–209 (1986). 10.1146/annurev.nu.06.070186.001143

5 Jacobsen, J. G. & Smith, L. H. Biochemistry and physiology of taurine and taurine derivatives. Physiol Rev 48, 424–511 (1968). 10.1152/physrev.1968.48.2.424

6 Shi, X., Yao, D. & Chen, C. Identification of N-acetyltaurine as a novel metabolite of ethanol through metabolomics-guided biochemical analysis. J Biol Chem 287, 6336–6349 (2012). 10.1074/jbc.M111.312199

7 Miyazaki, T. et al. Increased N-Acetyltaurine in the Skeletal Muscle After Endurance Exercise in Rat. Adv Exp Med Biol 975 Pt 1, 403–411 (2017). 10.1007/978-94-024-1079-2_33

8 Komine, S. et al. Taurine supplementation enhances endurance capacity by delaying blood glucose decline during prolonged exercise in rats. Amino Acids 54, 251–260 (2022). 10.1007/s00726-021-03110-8

9 Luginbühl, M., Rutjens, S., König, S., Furrer, J. & Weinmann, W. N-Acetyltaurine as a novel urinary ethanol marker in a drinking study. Anal Bioanal Chem 408, 7529–7536 (2016). 10.1007/s00216-016-9855-7

10 Meyre, D. et al. Genome-wide association study for early-onset and morbid adult obesity identifies three new risk loci in European populations. Nat Genet 41, 157–159 (2009). 10.1038/ng.301

11 Jong, C. J., Sandal, P. & Schaffer, S. W. The Role of Taurine in Mitochondria Health: More Than Just an Antioxidant. Molecules 26 (2021). 10.3390/molecules26164913

12 Jia, F. et al. Taurine is a potent activator of extrasynaptic GABA(A) receptors in the thalamus. J Neurosci 28, 106–115 (2008). 10.1523/JNEUROSCI.3996-07.2008

13 Huxtable, R. J. Physiological actions of taurine. Physiol Rev 72, 101–163 (1992). 10.1152/physrev.1992.72.1.101

14 Ito, T. et al. Taurine depletion caused by knocking out the taurine transporter gene leads to cardiomyopathy with cardiac atrophy. J Mol Cell Cardiol 44, 927–937 (2008). 10.1016/j.yjmcc.2008.03.001

15 Warskulat, U. et al. Taurine transporter knockout depletes muscle taurine levels and results in severe skeletal muscle impairment but leaves cardiac function uncompromised. FASEB J 18, 577–579 (2004). 10.1096/fj.03-0496fje

16 Ito, T., Yoshikawa, N., Schaffer, S. W. & Azuma, J. Tissue taurine depletion alters metabolic response to exercise and reduces running capacity in mice. J Amino Acids 2014, 964680 (2014). 10.1155/2014/964680

17 Warskulat, U. et al. Chronic liver disease is triggered by taurine transporter knockout in the mouse. FASEB J 20, 574–576 (2006). 10.1096/fj.05-5016fje

18 Waldron, M., Patterson, S. D., Tallent, J. & Jeffries, O. The Effects of an Oral Taurine Dose and Supplementation Period on Endurance Exercise Performance in Humans: A Meta-Analysis. Sports Med 48, 1247–1253 (2018). 10.1007/s40279-018-0896-2

19 Singh, P. et al. Taurine deficiency as a driver of aging. Science 380, eabn9257 (2023). 10.1126/science.abn9257

20 Park, E. et al. Cloning of murine cysteine sulfinic acid decarboxylase and its mRNA expression in murine tissues. Biochim Biophys Acta 1574, 403–406 (2002). 10.1016/s0167-4781(01)00364-5

21 McCoy, J. G. et al. Structure and mechanism of mouse cysteine dioxygenase. Proc Natl Acad Sci U S A 103, 3084–3089 (2006). 10.1073/pnas.0509262103

22 Veeravalli, S. et al. Flavin-Containing Monooxygenase 1 Catalyzes the Production of Taurine from Hypotaurine. Drug Metab Dispos 48, 378–385 (2020). 10.1124/dmd.119.089995

23 Dominy, J. E. et al. Discovery and characterization of a second mammalian thiol dioxygenase, cysteamine dioxygenase. J Biol Chem 282, 25189–25198 (2007). 10.1074/jbc.M703089200

24 Falany, C. N., Johnson, M. R., Barnes, S. & Diasio, R. B. Glycine and taurine conjugation of bile acids by a single enzyme. Molecular cloning and expression of human liver bile acid CoA:amino acid N-acyltransferase. J Biol Chem 269, 19375–19379 (1994).

25 Hasselmo, M. E. The role of acetylcholine in learning and memory. Curr Opin Neurobiol 16, 710–715 (2006). 10.1016/j.conb.2006.09.002

26 Grevengoed, T. J. et al. -acyl taurines are endogenous lipid messengers that improve glucose homeostasis. Proc Natl Acad Sci U S A 116, 24770–24778 (2019). 10.1073/pnas.1916288116

27 Chalhoub, G. et al. Carboxylesterase 2 proteins are efficient diglyceride and monoglyceride lipases possibly implicated in metabolic disease. J Lipid Res 62, 100075 (2021). 10.1016/j.jlr.2021.100075

28 Kim, J. T., Li, V. L., Terrell, S. M., Fischer, C. R. & Long, J. Z. Family-wide Annotation of Enzymatic Pathways by Parallel In Vivo Metabolomics. Cell Chem Biol 26, 1623–1629.e1623 (2019). 10.1016/j.chembiol.2019.09.009

29 Li, V. L. et al. An exercise-inducible metabolite that suppresses feeding and obesity. Nature 606, 785–790 (2022). 10.1038/s41586-022-04828-5

30 Jansen, R. S. et al. N-lactoyl-amino acids are ubiquitous metabolites that originate from CNDP2-mediated reverse proteolysis of lactate and amino acids. Proc Natl Acad Sci U S A 112, 6601–6606 (2015). 10.1073/pnas.1424638112

31 Varadi, M. et al. AlphaFold Protein Structure Database: massively expanding the structural coverage of protein-sequence space with high-accuracy models. Nucleic Acids Res 50, D439–D444 (2022). 10.1093/nar/gkab1061

32 Long, J. Z. et al. The Secreted Enzyme PM20D1 Regulates Lipidated Amino Acid Uncouplers of Mitochondria. Cell 166, 424–435 (2016). 10.1016/j.cell.2016.05.071

33 Cheng, A. G. et al. Design, construction, and in vivo augmentation of a complex gut microbiome. Cell 185, 3617–3636.e3619 (2022). 10.1016/j.cell.2022.08.003

34 Preising, M. N. et al. Biallelic mutation of human. FASEB J 33, 11507–11527 (2019). 10.1096/fj.201900914RR

35 Chen, C. et al. Roles of taurine in cognitive function of physiology, pathologies and toxication. Life Sci 231, 116584 (2019). 10.1016/j.lfs.2019.116584

36 Emmerson, P. J. et al. The metabolic effects of GDF15 are mediated by the orphan receptor GFRAL. Nat Med 23, 1215–1219 (2017). 10.1038/nm.4393

37 Dodd, D. et al. A gut bacterial pathway metabolizes aromatic amino acids into nine circulating metabolites. Nature 551, 648–652 (2017). 10.1038/nature24661

